# A quantitative pipeline for whole-mount deep imaging and analysis of multi-layered organoids across scales

**DOI:** 10.1101/2024.08.13.607832

**Authors:** Alice Gros, Jules Vanaret, Valentin Dunsing-Eichenauer, Agathe Rostan, Philippe Roudot, Pierre-François Lenne, Léo Guignard, Sham Tlili

**Author notes:** These authors contributed equally as co-first authors (Alice Gros, Jules Vanaret, Valentin Dunsing-Eichenauer). The order of authors reflects the relative amount of time each author dedicated to the project. These authors contributed equally as co-last authors (Pierre-François Lenne, Léo Guignard, Sham Tlili).

## Abstract

Whole-mount 3D imaging at the cellular scale is a powerful tool for exploring complex processes during morphogenesis. In organoids, it allows examining tissue architecture, cell types, and morphology simultaneously in 3D models. However, cell packing in multilayered organoid tissues hinders both deep imaging and quantification of cell-scale processes. To address these challenges, we developed an experimental and computational pipeline to extract properties at scales ranging from cell to tissue. The experimental module is based on two-photon imaging of immunostained organoids. The computational module corrects for optical artifacts, performs accurate 3D nuclei segmentation and reliably quantifies gene expression. We provide the computational module as a user-friendly Python package called Tapenade, along with napari plugins which enable joint data processing and exploration across scales. We demonstrate the pipeline by quantifying 3D spatial patterns of gene expression and nuclear morphology in gastruloids, revealing how local cell deformations and gene co-expression relate to tissue-scale organization. This quantitative pipeline improves our understanding of gastruloid development, and lays the groundwork for a wide range of multi-layered organoids and tumoroids systems

## INTRODUCTION

### Understanding organoid development and variability at multiple scales

The biology of organoids and tumoroids represents a frontier in fundamental biology and biomedical research, driven by the need for accurate 3D models complementary to 2D cell cultures and animal studies [1, 2]. Tin the context of the “3Rs” (Replacement, Reduction, and Refinement) aimsed at minimizing the use of animals in research and enhancing ethical experimental practices [3]. Developing these models in 3D is crucial for mimicking the complex architecture and functionality of tissues, enabling deeper insights into developmental or disease mechanisms and treatment responses [4, 5]. Organoids models exhibit less stereotypic development compared to embryos, where cell identification is typically straightforward based on position within the embryo at specific developmental stages [6, 7]. In contrast, organoids present more complex developmental trajectories [8–11], characterized by variability in morphology, cell type composition, and differentiation levels. This variability may stem from differences in the initial cell differentiation state or number [12], as well as from a less controlled biochemical and mechanical environment than that of embryos [13], which grow confined within maternal tissues for mammals or in a rigid shell for insects [14]. This inherent variability underscores the need for detailed characterization of a sufficient number of organoids in parallel to properly characterize the diversity of their developmental processes. To this end, coarse-grained methods have been developed to classify phenotypes of intestinal organoids [15] and gastruloids [16, 17] by analyzing maximum projections from 3D immunofluorescence staining using high-throughput confocal imaging. These approaches are effective for systematically screening the impact of chemical compounds or signaling pathway perturbations on organoid morphology, as well as on the localization and proportion of various cell types and tissues within the organoid.

However, understanding how organoids or tumoroids develop and self-organize also requires capturing how individual stem cells or cancerous cells differentiate and modulate their behavior and morphology in response to their local microenvironment [18, 19]. Simultaneously, it is essential to quantify tissue characteristics on a coarser scale (where each measurement integrates data from groups of several neighboring cells) to capture the mechanical and biochemical interactions among different tissues and cell populations within the overall geometry of the organoid [20]. To achieve this, a challenge is to image these 3D tissues *in toto* at a resolution sufficient to identify each cell and its neighbors both in live and fixed samples [21]. On fixed samples, multiple specific fluorescent markers can be used to capture cell fate by measuring the expression levels of particular genes or proteins by immunostaining or in situ hybridization. Complementary, live imaging is essential for understanding how local cellular events (division, migration, rearrangements) underlie the morphogenesis of these systems, as it has been done previously for embryonic development [22, 23].

### Imaging organoids *in toto* at cell resolution

Recently, light-sheet and confocal imaging systems have been optimized for parallel imaging or long-term live imaging of organoids [24–27]. Using deep-learning-based segmentation, tracking, and classification tools, these protocols generate digital atlases of developing organoids at single-cell resolution. For example, [25] employed a dual illumination inverted light-sheet microscope to live-image gut organoids over several days, enabling cell lineage reconstruction from nuclei tracking. In [26], a single-objective light-sheet microscope was used to image tens of 3D tissues, such as gut, hepatocyte, and neuroectoderm organoids, in parallel for phenotypic classification.

While advances in light-sheet microscopy have extended imaging depth in organoids, maintaining high image quality throughout thick samples remains challenging. In practice, quantitative analyses are still largely restricted to organoids under roughly 100 µm in diameter [24, 28, 29], or hollow organoids with cavities, lumens, and epithelial structures [25]. These organoids are less optically opaque than larger organoids like gastruloids [30, 31], neuromuscular organoids [32], or cancer spheroids [33], which can reach diameters of 300 microns and more.

For large organoids, multiphoton microscopy provides a powerful alternative due to its ability to penetrate deep into thick tissues with minimal photodamage [34]. This technique utilizes longer wavelengths of light to excite fluorescent molecules within the specimen, allowing the visualization of cellular structures and interactions in high resolution [35]. Furthermore, it avoids drawbacks of confocal or light-sheet microscopy on large, densely packed samples, such as strong intensity gradients, image blurring, and reduced axial information due to light scattering, aberrations, degradation or divergence of the light-sheet [36]. Finally, two-photon microscopes are typically more accessible than light-sheet systems and allow for straightforward sample mounting, as they rely on procedures comparable to standard confocal imaging.

### Gastruloids: a toy model for an integrated pipeline to analyze organoids development *in toto* at cell resolution

We have recently developed two-photon live imaging protocols to image developing gastruloids over multiple hours [31, 37]. Gastruloids are mouse embryonic stem cells that self-organize into 3D embryonic organoids, sharing similarities with their *in vivo* counterparts [30]. Recent studies have shown that, within a few days, gastruloids undergo significant morphological changes, developing structures that closely resemble organs both genetically and morphologically, such as neural tube-, [38] gut-, and cardiac-like structures [39]. Live two-photon imaging of gastruloids around their mid-plane has enabled us to propose biophysical models for their early symmetry breaking and elongation, using coarse-grained 2D analysis of collective tissue flows and gene patterning [37]. However, investigating in a full 3D context the relation between cellular fate and local tissue architecture during gastruloids morphogenesis [40] is still a key challenge. Indeed, it necessitates imaging *in toto* these dense 3D cell aggregates, which have a typical diameter above 200 microns and are highly light-diffusive objects.

In this work, we present an integrated pipeline to: (1) image multiple immunostained and cleared gastruloids in whole-mount configuration using two-photon microscopy, capturing both *in toto* and cellular scale details; (2) process and filter the resulting 3D images to correct optical artifacts from deep imaging and segment individual cell nuclei; and analyze gene expression patterns, cellular events, and morphologies in 3D across multiple spatial scales (Fig.1).

**Fig. 1.**
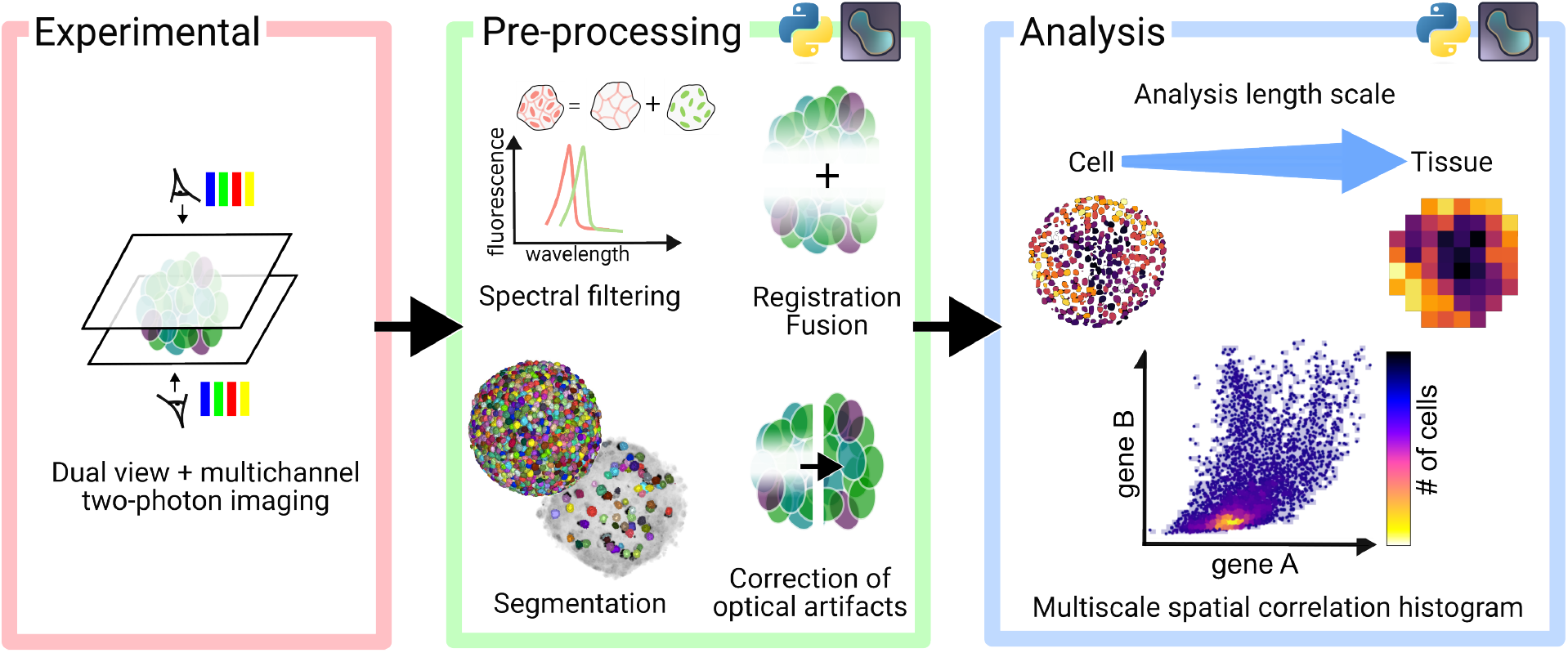
Overview of the experimental and computational pipeline for *in toto* imaging of organoids at cellular resolution and multiscale analyses.

## I. RESULTS

### An experimental and computational pipeline for *in toto* organoid imaging

We have developed a versatile pipeline to image, process and analyze tens of immunostained gastruloids (with diameters ranging from 100 to 500 *µm*). The experimental pipeline consists of sequential opposite-view multi-channel imaging of cleared samples, imaged with a commercial two-photon microscope (Fig.1). Afterwards, multiple processing steps are performed to extract meaningful information from the acquired images: i) spectral unmixing to remove signal cross-talk, ii) dual-view registration and fusion to reconstruct *in toto* images, iii) sample and single cell segmentation, and iv) signal normalization across depth and channels. Finally, reconstructed *in toto* images are analyzed at different scales, from cell-level correlations of different genes to coarse-grained maps of gene expression patterns, cell shapes, densities, and division events. All processing and analysis tools are open-source and Python-based. They are openly accessible as notebooks and interactive napari plugins.

### *In toto* multi-color two-photon imaging

To facilitate dual-view imaging, we mounted immunostained gastruloids between two glass coverslips using spacers of defined thickness (typically 250 − 500 *µm*, adapted to match the size of the gastruloids without compressing them), allowing us to image the sample iteratively from two opposing sides (Fig.2a, see Methods III A 3). To optimize imaging performance, we compared several refractive index matching mounting mediums, glycerol [41], ProLong Gold Antifade mounting medium (ThermoFisher), and optiprep [42], on gastruloids labeled with Hoechst nuclei stain (see Methods III A 1). We identified 80% glycerol as the mounting medium with best clearing performance (Fig.2b, Fig.S1c), leading to a 3-fold/ 8-fold reduction in intensity decay at 100 *µm*/ 200 *µm* depth compared to mounting in phosphate-buffered saline (PBS) and superior performance to gold antifade and live-cell compatible optiprep medium (Fig.S1c). At these depths, information content was significantly improved by 1.5- and 3-fold, quantified using Fourier ring correlation quality estimate (FRC-QE) [43] (Fig.2c). Segmenting cell nuclei in images acquired on glycerol-cleared samples reliably detected cells at depth up to 200 *µm*, whereas a continuous decline in cell density was observed for PBS mounting, with four times fewer cells detected at 200 *µm* depth (Fig.2c, see details on the segmentation in the next section). These degradation effects were much more pronounced in confocal imaging even in glycerol clearing, which showed 2× lower intensity and 8× lower FRC-QE than two-photon imaging at 100 *µm* depth (Fig.S1a,b).

**Fig. 2.**
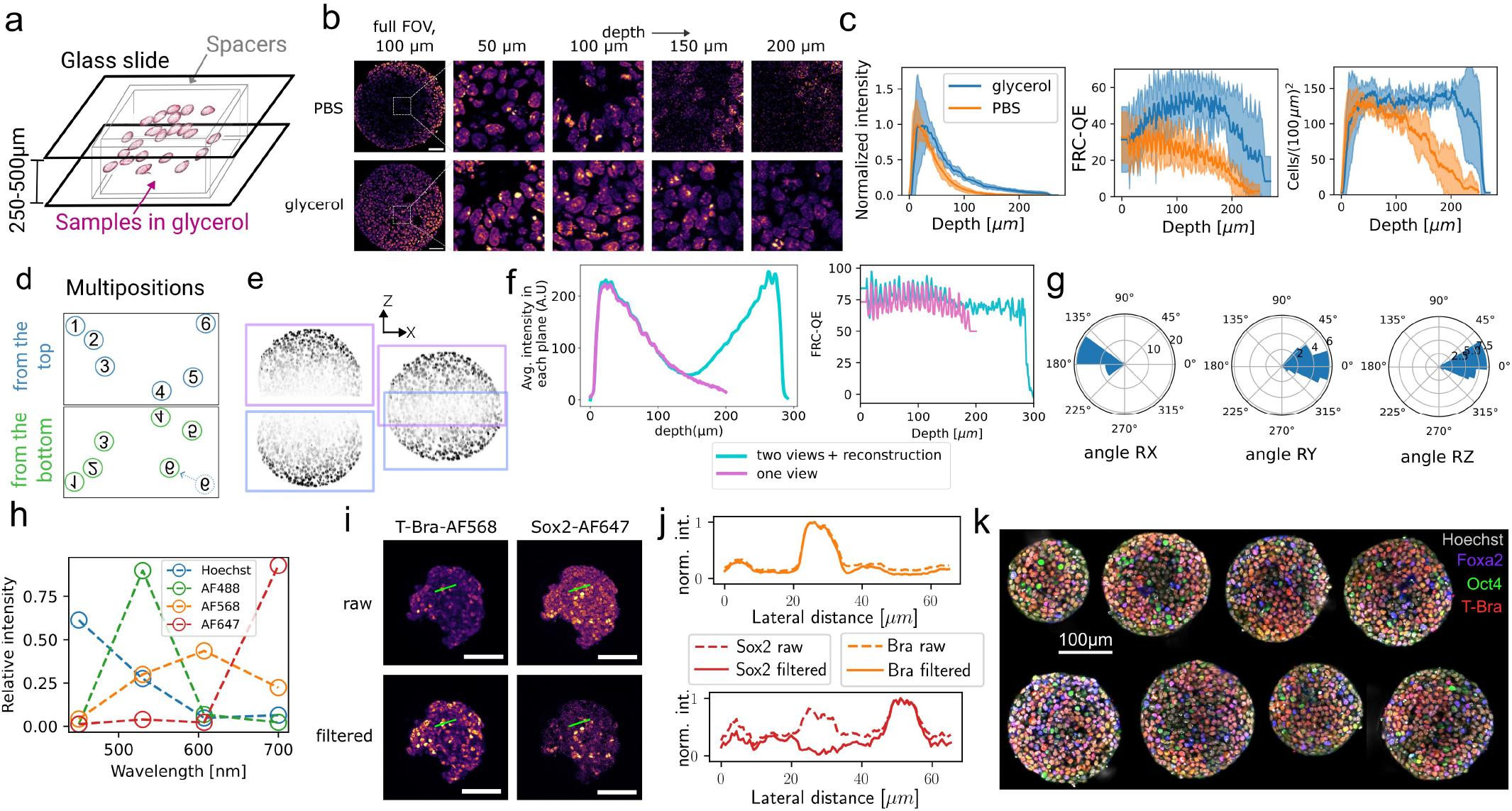
Dual-view multi-color two-photon imaging. **(a)** Multiple gastruloids are mounted in glycerol clearing medium in between two glass slides using spacers. **(b)** Mounting gastruloids in 80% glycerol significantly improves imaging at depth compared to PBS mounting. Insets show images at different depth in a central square region of 60 × 60 *µm*^2^. **(c)** Quantification of intensity, FRC-QE image quality metric and density of segmented nuclei as a function of depth in the central region depicted in (b) for both glycerol- and PBS-mounted samples. Shaded areas show standard deviation for six (PBS) and nine (glycerol) gastruloids. **(d-f)** Top and bottom positions are assigned using a pattern matching algorithm that is robust to residual movements of individual gastruloids, as illustrated for position 6 (d). Respective image stacks are registered using rigid 3D transformations and fused applying sigmoid fusion weights (e). This leads to *in toto* coverage of whole gastruloids, in which individual cells are still visible in the mid-plane (e), as shown for a gastruloid of typical size of 250 *µm* diameter. The average intensity per plane and FRC-QE are shown in (f) for one- and dual-view images. **(g)** Polar histogram plots of rotation angles around X-,Y-, and Z-axis from the 3D registration of dual-view top and bottom acquisitions, pooled for 37 aggregates. The sample slide was flipped around the X-axis. **(h-j)** Multi-color detection using two-photon excitation can lead to significant spectral cross-talk, as shown in the apparent emission spectra detected on gastruloids stained with a single fluorophore species (h). Using spectral unmixing, false-positive signal due to spectral cross-talk is effectively removed, as shown in the raw and filtered images (here at 50 *µm* depth) of gastruloids stained with T-Bra-AF568 and Sox2-AF647 (i) and the quantification of the intensity along a line (10 pixels width) across few nuclei for raw and spectrally filtered images (j). **(k)** Four-color images of the mid-plane of 77 h gastruloids stained with Hoechst, FoxA2-AF488, Oct4-AF568 and T-Bra-AF647, obtained with dual-view four-channel two-photon imaging and spectral unmixing. The gastruloids were acquired in the same experiment. Scale bars are 50 *µm* in (b,i) and 100 *µm* in (k).

Nevertheless, two-photon microscopy in glycerol beyond 200 *µm* depth suffered from decreased signal-to-noise-ratio (SNR) (Fig.2b,c). To properly image larger gastruloids *in toto*, a second detection side was required. We, therefore, flipped the sample slide and re-imaged the same set of gastruloids from the opposing side. Whereas the flipping was done manually, all mounted gastruloids were imaged automatically after defining their positions in the imaging software. A typical acquisition took about 5-10 min per gastruloid per side for 1 *µm* z-spacing and full field-of-view (318 *µm* with 0.62 *µm* pixel size at 40× magnification). To reconstruct *in toto* image stacks, we automatically identified opposite views of the same gastruloid by applying a pattern matching algorithm to gastruloids positions in the two acquisitions (Fig.2d, see Methods III F). Next, we registered each pair of image stacks acquired on the same aggregate using a rigid 3D transformation determined by a content-based block-matching registration algorithm previously applied for multi-view and time registration of light-sheet imaging data [6, 7]. Finally, the registered image stacks were fused using a sigmoid decay function to weight contributions of the two sides (Fig.2e,f and S5). The registration was performed on the ubiquitous nuclei channel, i.e. Hoechst staining. In some instances, flipping of the slide resulted in additional minor rotations (on average (− 13 ± 11)^*°*^, (3 ± 18)^*°*^ and (5 ± 10)^*°*^ around the X-, Y-, and Z-axis for n=37 aggregates), which occasionally prohibited convergence of the registration algorithm (Fig.2g). For these cases, we implemented a napari plugin to pre-register the images manually and visualize the result in 3D (Fig.6). Fused gastruloid images exhibit about 4-fold lower intensity in the mid-plane compared to the outer ends, but approximately constant FRC-QE across depth (Fig.2f).

A common limitation in fluorescence imaging is the low number of targets that can be excited simultaneously and spectrally discriminated. To image multiple targets in parallel, we excited four fluorophore species (Hoechst, AF488-, AF568-, AF647-tagged secondary antibodies on immunostained gastruloids) simultaneously. Thanks to the broad two-photon excitation spectra, the four dye species could be excited with just two laser lines, 920 nm and 1040 nm. However, detecting their emission simultaneously on four non-descanned detectors resulted in significant signal cross-talk, due to spectral overlap (Fig.2h-j). We circumvented this problem by applying spectral decomposition [45]. We calibrated spectral patterns, i.e. apparent emission spectra for each separate fluorophore species, in gastruloids stained with a single fluorophore species, and used these as reference patterns to decompose multi-color data (Fig.2h). Because of chromatic effects due to dispersion, spectral patterns were not constant at different depths in the sample. For example, shorter wavelength emission decays stronger with imaging depth than longer wavelength fluorescence (Fig.S3b). To account for this, we performed the unmixing with depth-dependent spectral patterns (see Methods III E). Thereby, we could significantly remove cross-talk, for example, false positive cells in the far-red detection channel that resulted from cross-talk of bright cells visible in the red channel (Fig.2i,j). Notably, spectral decomposition with four-channel detection was even possible for spectrally strongly overlapping fluorophores, which we verified on gastruloids with transgenic H2B-GFP expression in the nuclei, stained with E-cadherin (Ecad)-AF488 at cell membranes (Fig.S1e,f). We thus applied spectral unmixing to all data presented throughout this work. This way, we could generate cross-talk free four-color images of several biologically relevant markers imaged simultaneously in multiple gastruloids in the same experiment (Fig.2k, Suppl.Movies 1-3).

### Single cell segmentation

Quantifying cell-scale gene expression and nuclei morphological properties requires accurate instance segmentation of individual nuclei. In early-stage gastruloid datasets acquired with two-photon microscopy, accurate segmentation of nuclei is hindered by high nuclei packing (Fig.2b), shape heterogeneity, and spatially varying SNR due to optical aberrations and scattering, especially deeper in the sample. We qualitatively assessed the performance of two open-source state-of-the-art methods for 3D segmentation of nuclei in dense environments, i.e Cellos [46] and AnyStar [47], and observed unsatisfying results (Fig.S9a). We also considered 3DCellScope [29], but its closed-source nature and required a paid licence to access full segmentation features. We thus trained a custom Stardist3D [48] model to segment nuclei, as it excels with separating star-convex objects in close contact.

A mid-plane section of the StarDist prediction is shown Fig.3a, exhibiting a qualitatively good result on outer layers as well as inner part of the sample. To increase robustness, the network was trained on three datasets acquired with two-photon microscopy in different experimental conditions, namely on (1) live-imaging of Histone 2B (H2B)-GFP gastruloids, (2) live-imaging of sparsely labeled mosaic gastruloids (composed of a mix of 75% of non-fluorescently labeled cells and 25% of cells expressing H2B-GFP under the Brachyury gene promoter [31]), and (3) 96 h fixed samples stained with Hoechst. This combined dataset presented variations in labeled nuclei density, nuclei texture, and in voxel dimensions (Fig.3b, see table II and Fig.S2a). In total, we annotated 4414 nuclei in 3D and used data augmentation (see Methods III G) to enrich the training datasets (Fig.3b). Both during training and inference, we applied local contrast enhancement and normalization to all datasets by clipping intensity histograms computed in local boxes at the 1st and 99th percentile and by mapping intensities between 0 and 1. This step was crucial to homogenize the intensity distribution across and inside datasets during training, and improved segmentation quality by reducing the number of false detections at depths at which the SNR is reduced.

**Fig. 3.**
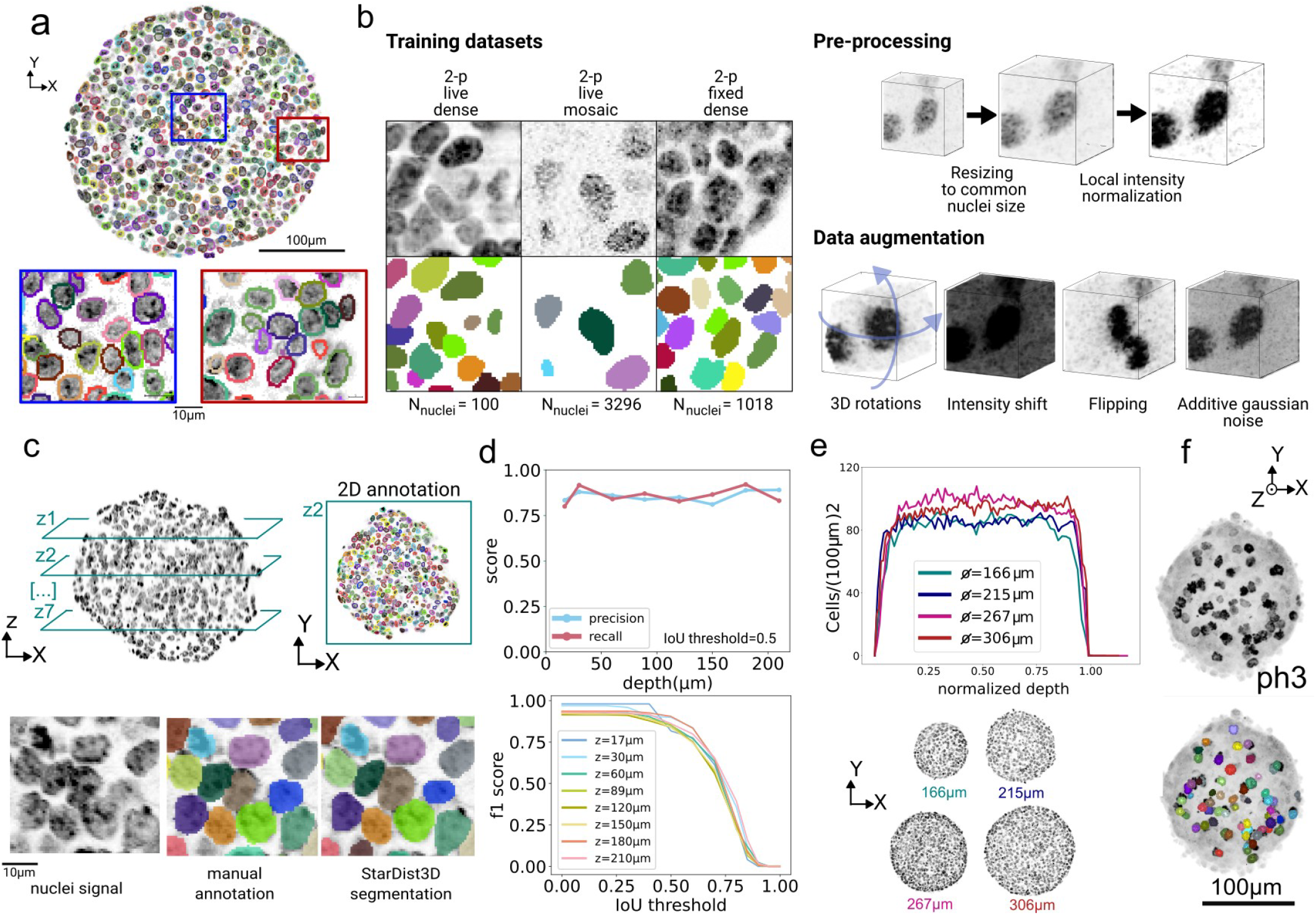
3D nuclei segmentation with StarDist3D. **(a)** Exemplary segmentation (contours) of Hoechst stained nuclei (gray) using custom trained StarDist3D model. Insets show segmentation results in inner and outer cell layers, at 150 *µm* depth. **(b)** Three different datasets were annotated and used for training. Images show exemplary annotated patches (top row: intensity images, bottom row: label images). Before training, images were resampled to a common voxel size of 0.62 × 0.62 × 0.62 *µm*^3^. Image contrast when enhanced using local histogram clipping and normalization. During training, data augmentation was used to maximize training input. **(c)** To evaluate segmentation performance, entire z-planes of a gastruloid were annotated at different depths in 2D, on a 250 *µm* size gastruloid imaged from two views and fused, and the segmentation result was compared with the ground truth. **(d)** Top : Precision and Recall at IoU threshold 0.5 as a function of depth. Bottom: F1 score as a function of IoU thresholds, color represents depth. **(e)** Cell density obtained from 3D segmentation of gastruloids of different diameters. The corresponding images at the mid-plane are shown at the bottom. **(f)** 3D rendering of the detection of mitotic cells using StarDist3D. Top image shows ph3 stained sample, bottom image shows detected cell divisions overlayed with the ph3 signal.

To assess the performance of the training, we manually annotated 2D planes at different depths throughout a sample (see Fig.S2c, sample 1) and compared the result of StarDist3D to the ground truth, both in terms of quality of nuclei detection and volume reconstruction (Fig.3c). To evaluate the performance, we computed the recall and the precision (resp. quantifying how many relevant items are retrieved and how many retrieved items are relevant), and the F1 score as the harmonic average between the two. We show in Fig.3d that our trained model recovers a constant precision and recall across the full depth, and the F1 score is superior to 0.8 up to intersection over union (IoU) threshold 0.5. Comparing the recall and precision to the ones obtained with global contrast enhancement, i.e., histogram clipping based on the percentiles of the whole image, our local contrast enhancement scheme mitigates both false positives and false negatives in deeper z planes (Fig.S2d-e). It is still necessary to image from two views to recover all the nuclei, as shown Fig S2f-g, even using the local contrast enhancement method. Our score measurement is based on computing the IoU of 3D objects based on 2D images. As manually annotating many dense organoid datasets for validation is a slow and cumbersome process, we designed a validation method based on comparing the 3D prediction of StarDist3D with 2D annotations in several XY planes along the z-axis. We validate this approach on Fig.S2b using four different samples in which we chose XY planes spaced evenly along the z-axis, and on which nuclei were annotated in 3D. The F1 score obtained in 3D (3D prediction compared with the 3D ground-truth) is well approximated by the F1 score obtained in 2D (2D predictions compared with the 2D sliced annotated segments). The difference between the 2 scores was at most 5%. Finally, we observed that the model preserves a constant cell density across depth in spherical gastruloids of up to 475 *µm* in diameter (Fig.3e).

Overall, our custom model achieved an F1 score of 85 ± 3% at 50% IoU. Using a variety of annotated datasets, dualview imaging and a local contrast enhancement algorithm, we trained a powerful StarDist3D model that shows robust segmentation performances regardless of depth, sample size, and experimental conditions.

### Maps of proliferation and cell density

Differential division and growth rates in tissues are key drivers of morphogenetic changes [49]. Therefore, quantifying tissue-scale gradients in cell proliferation is crucial for understanding the morphogenesis of gastruloids. To detect division events, we stained gastruloids with phosphohistone H3 (ph3) and trained a separate custom Stardist3D model using 3D annotations of nuclei expressing ph3 (see Methods III H). This model together allowed us to detect nearly all mitotic nuclei in whole-mount samples for any stage and size (Fig.3f and Suppl.Movie 4), and we used minimal manual curation to correct remaining errors. To probe quantities related to the tissue structure at multiple scales, we smooth their signal with a Gaussian kernel of width *σ*, with *σ* defined as the spatial scale of interest. From the segmented nuclei instances, we compute 3D fields of cell density (number of cells per unit volume), nuclear volume fraction (ratio of space occupied by nuclear volume within the local averaging volume, as defined in the Methods III I with a scale of *σ*), and nuclear volume at multiple scales. We plot each field on Fig.4a at three spatial scales ranging from the nuclei scale, i.e *σ* set to the average nuclear radius, to the tissue scale, i.e *σ* set to six times the average nuclei radius. Taking this smoothing approach, we computed fields of division density (number of division events per unit volume), that represent the local density of cells in mitosis, from the division labels detected on ph3 as described above. An increased volumetric division density can result from either elevated cell proliferation or higher cellular volumetric density. To distinguish between these effects, we computed a proliferation probability field that represents the local fraction of cells undergoing mitosis within the 3D sample (by dividing the division density field by the cell density field). A cross-section of each of these 3D maps is represented in Fig.4b, showing a typical heterogeneous pattern of division density and cell density. These maps can be visualized in parallel for several samples, which allows systematic comparison of patterns for high-throughput acquisitions. Analyzing proliferation fields in 3D gives us insight on growth patterns during development and is key to understand and quantify gastruloid morphogenesis. Importantly, this quantification can be applied across various developmental stages, providing quantitative insights into growth phases under different stages or media conditions. A proliferation pattern for a strongly elongated 120 h gastruloid grown in a modified culture protocol is shown in Fig.S6b.

**Fig. 4.**
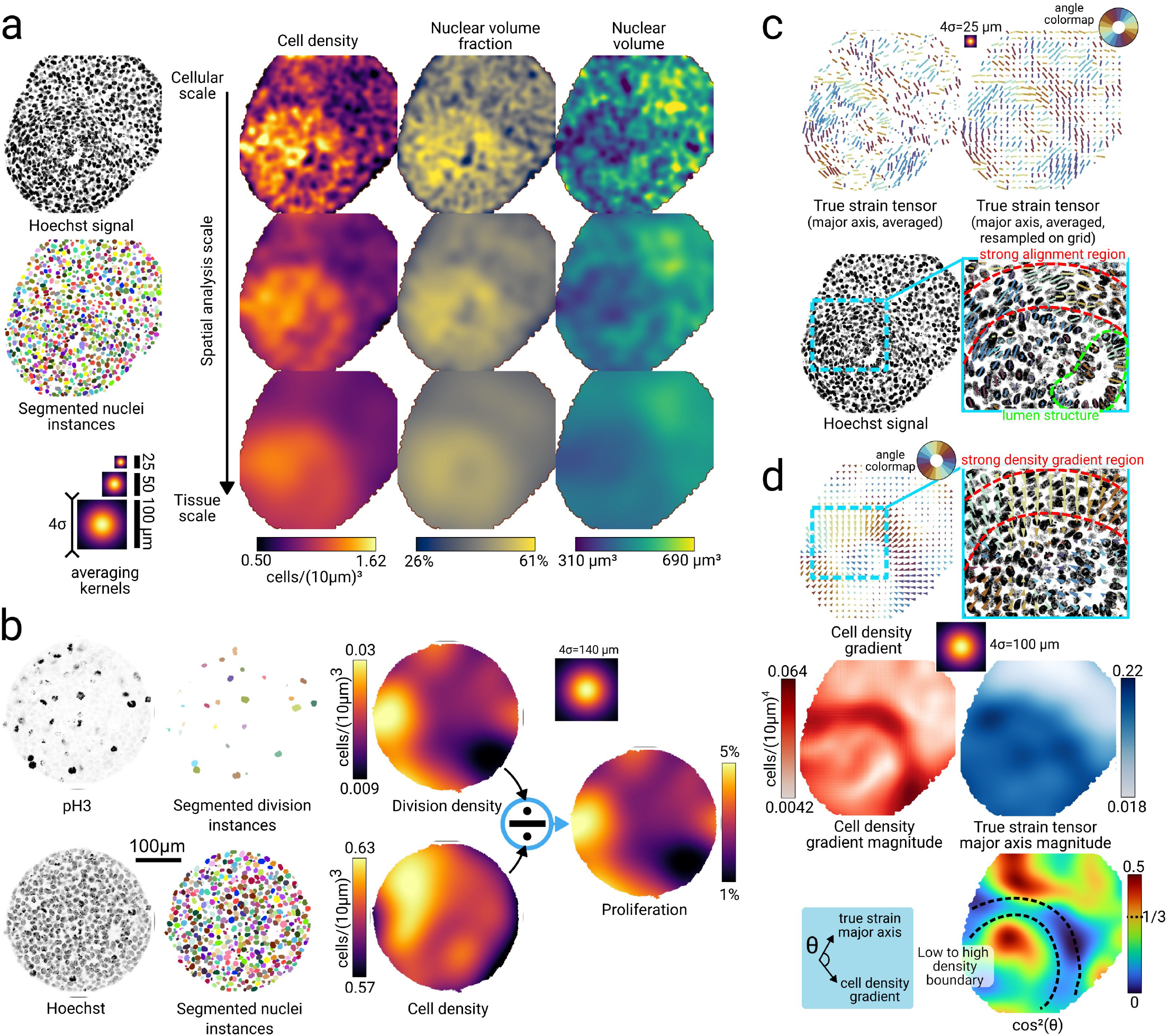
Cell-to-tissue scale morphological analysis. **(a)** Segmenting individual nuclei instances in 3D allows a complete characterization of object packing by quantifying cell density, nuclear volume fraction, and nuclear volume. Each quantity is shown at three resolutions ranging from the cell scale to the tissue scale, which correspond to convolution with Gaussian averaging kernels with *σ* = 6, 12, and 25 *µm*, respectively. **(b)** Divisions stained with ph3 and nuclei stained with Hoechst are segmented using StardDist3D and used to compute respectively maps of division density and cell density. The proliferation is defined as the ratio of the division density with the cell density, showing the local fraction of cells undergoing mitosis. **(c)** Cell-scale fields of deformation, quantified by the major axis of the nuclei true strain tensors. A version sampled on a regular grid is provided for better readability. The deformation field reveals a region of spatially persistent alignment that overlaps with the density boundary and highlights nuclei forming the lining of a lumen. **(d)** We compute the tissue-scale field of squared cosine of the angle between the cell density gradient and the true strain tensor major axis to quantify local regions of anti-alignment. The vector magnitudes of the two fields are shown to add perspective to the analysis in regions where the gradient or the local deformation is small. All quantities shown in this figure are computed in 3D, but we show a single z-slice (vector quantities are projected onto the slice) for convenience.

### Morphometric analysis

Tissue-scale patterns of cell/nuclei deformations in developing embryos or organoids are good indicators of the existence of mechanical stresses which can contribute to tissue morphogenesis [50] and affect in return cell differentiation [51]. To explore changes in tissue structure and organization during gastruloid development, we studied spatial heterogeneities in cell densities and cell deformations emerging during the first morphogenetic phase of gastruloids development, which consists in a global tissue elongation between 72 and 96 h after their initial aggregation. We have shown in previous work [31, 37] that such elongation is associated with the polarization of the gastruloid, exhibiting a posterior pole rich in E-cadherin and constituted of a mixed population of both endoderm and nascent mesoderm tissues, and an anterior pole poor in E-cadherin constituted of cardiac and vascular mesoderm tissues which are mesenchymal. Here, to prevent any relaxation of cellular deformation caused by tissue fixation, we performed 3D acquisition of live gastruloids with an H2B-GFP marker. We illustrate the full approach on a single sample in Fig.4c-d, and show that the findings are reproducible across different samples and views (see Fig.S6a). The cell density field revealed a high level of heterogeneity at the nuclei scale, but analysis at different scales always showed the presence of two tissue-scale populations of low and high cell densities located respectively in the posterior and anterior part of the sample, separated by a sharp boundary characterized by the length scale of a few nuclei. These two populations and the boundary were also visible, although less clearly, from the nuclear volume and nuclear volume fraction fields, with the high-density population showing a higher nuclear volume fraction but lower nuclear volumes. The nuclear volume fraction indicates regions of high packing located in the high-density population, with a maximum of 61%, close to the theoretical value for random close packing of spheres (64%). As extra-cellular space volume fraction is low in such sample [37], this indicates high nuclei-to-cell volume ratios, which suggests that nuclei shapes are a good proxy for cell shapes, and thus nuclei could be relevant in cell-scale mechanotransduction processes.

To quantify cell deformation, we first computed the inertia tensor from each segmented nuclei instance and obtained the true strain tensor [52] for each cell (see Methods III L), which is a size independent measure of object deformation. This justifies averaging these tensors among nuclei with large size variations by effectively removing the size bias. We plot the major axis (associated with the largest eigenvalue) of the true strain tensor for each nucleus on Fig.4 at the nuclei scale. The major axis is oriented along the direction of the largest deformation of the nuclei. Close to the low-to-high density boundary, we observed that nuclei are elongated in the direction of the boundary, i.e perpendicular to the gradient of cell density. We further quantified this observation by computing the angle *θ* between the gradient of cell density and the major axis of the true strain tensors. In Fig.4d, we plot the continuous field representing the quantity cos^2^ (*θ*) averaged at the tissue scale. For random orientations of the major axis with respect to the gradient of cell density, the average value of cos^2^ (*θ*) is 1*/*3, and the value approaches 0 or 1 for respectively perpendicular and parallel orientations. The field reveals a wide region with values below 0.1 around the boundary, confirming that nuclei are elongated along the tissue boundary. Other regions show comparatively smaller values of the local gradient amplitude or no tissue-scale alignment. These descriptions appear reproducibly in several samples, see Fig.S6a, and are consistent with an interpretation of the boundary as a region where mechanical stresses are generated between gastruloids anterior and posterior poles. This analysis also shows how nuclei shapes at small scales are appropriate proxies for deformation inside and around tissue micro-structures like lumen and cavities, which allows for an accurate description of the local epithelial architecture. At the tissue scale, nuclei shapes can serve as a proxy for tissue mechanical stresses in the context of tissues modeled as visco-elastic liquids with residual elasticity in cell deformation [53].

### Quantitative analysis of gene expression at global and cell-scale level

A key interest in *in toto* imaging is that it allows quantifying expression gradients of genetic markers and proteins of interest across entire organoids. In the case of gastruloids, gradients of differentiation emerge during Wnt activation, prior or concomitant with symmetry breaking [16, 31, 54, 55]. Due to the previously mentioned limitations of light-sheet microscopy for *in toto* coverage and quantitative intensity analyses for gastruloids beyond 100 − 200 *µm* diameter, investigations of such gradients have thus far been restricted to analyses of low resolution 2D images [16, 54]. Because of this limitation, it is still unknown when gradients of differentiation are established during gastruloid development. To explore this, we analyzed images of gastruloids fixed at 77 h, labeled with Hoechst and additionally immunostained with T-Bra, a mesendoderm marker expressed upon Wnt mediated cell differentiation. Previous investigations with 2D imaging indicated a gradient of higher T-Bra expression in outer cell layers and lower expression in the interior [31].

Although our dual-view imaging pipeline delivers *in toto* coverage, images of the ubiquitous Hoechst nuclei stain still exhibited intensity gradients along the z direction and between outer and inner cell layers (Fig.5a). We attributed these gradients to optically induced heterogeneities and developed a correction method that re-normalizes the channels of interest based on the local nuclei intensity. To this aim, we first corrected the intensity decrease, which is dependent on the emission wavelength and the depth (see Fig.S3). We applied on this corrected signal a convolution with a 3D Gaussian kernel masked with the nuclei segmentation. The result of that convolution is a coarse-grained map of nuclei intensity, that we use to locally normalize the original image (Fig.5a). Since this normalization was done in 3D, the intensity values in the mid-plane of a gastruloid were enhanced while the intensities in the first and last z-planes remained constant (Fig.S8b). Furthermore, the method effectively and robustly homogenized nuclei intensities across gastruloids from different acquisitions and imaging conditions (Fig.S8c and d). We evaluated different sizes for the Gaussian kernel of the convolution and chose an optimal one, with a typical value around the diameter of the nuclei (10 − 15 *µm*), that lead to the most homogeneous Hoechst signal after re-normalization (see Methods III J and Fig.S8e). This normalization scheme accounts for heterogeneities in gastruloids of arbitrary shape as well as local cell density and thus variations in the optical paths of excitation and fluorescence light rays that cannot be captured by a simple geometrical model.

**Fig. 5.**
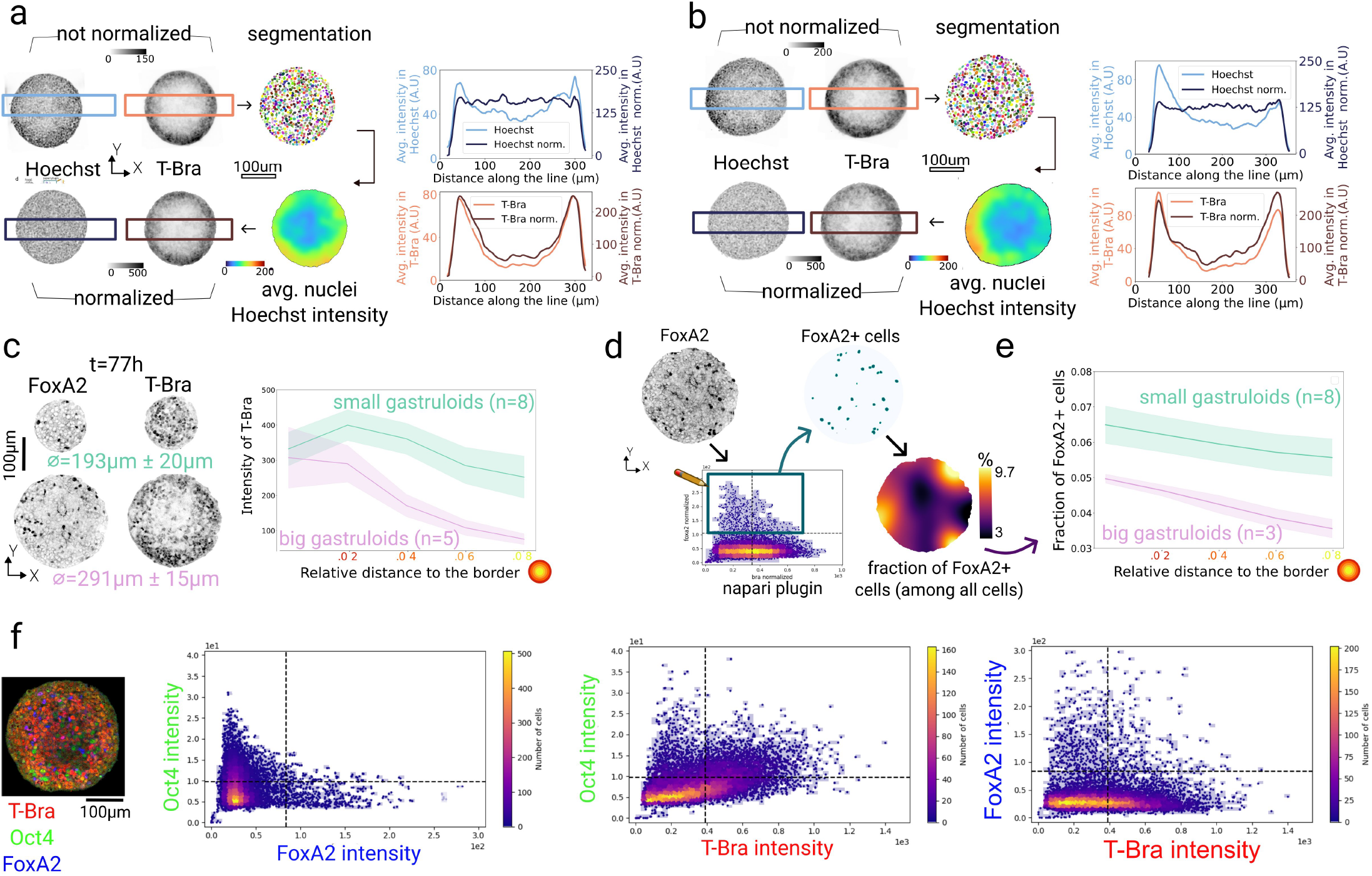
Coarse-grained and cell-level gene expression analysis. **(a,b)** Images of Hoechst and T-Bra in the mid-plane of 77 h gastruloids. Using the nuclei segmentation, maps of averaged nuclei intensity were generated and used to locally re-normalize the T-Bra signal. The graphs on the right side of each panel show the intensity across rectangular sections of the mid-plane, shown in the images as blue/ brown boxes, before and after intensity normalization. This intensity profile is smoothed using a Gaussian kernel on the images to extract the large-scale intensity variations. (a) example of a radially symmetric gradient of nuclei intensity, from the exterior to the interior. (b) example of an asymmetric gradient, the left side of the sample has a high nuclei intensity that is corrected after normalization, changing the T-Bra profile (bottom right graph). **(c)** Images of 77 h gastruloids (mid-plane) of different size, immunostained for T-Bra and FoxA2 gene expression markers. From the re-normalized 3D images, radial intensity plots were generated for big (291 ± 15 *µm* average diameter, n=5) and small (193 ± 20 *µm* average diameter, n=8) gastruloids as a function of distance to the border. Shaded regions show standard deviations. **(d)** For one of the previously described samples fixed at 77 h, FoxA2 positive cells are visualized from the renormalized data using the interactive napari tools and the image thresholded accordingly. Using the binary image of FoxA2 positive cells, a map of local fraction of FoxA2 positive cells, rescaled by the local cell density, was computed, showing a radial pattern. **(e)** Analysis of the radial distribution, i.e. fraction of FoxA2 positive cells as a function of distance to the border, for big (278 ± 16 *µm* average diameter, n=3) and small (193 ± 20 *µm* average diameter, n=8) aggregates, computed from the coarse-grained maps as shown in (d). Shaded regions show standard deviations. **(f)** Three-color image of a 77 h gastruloid immunostained for T-Bra, Oct4, and FoxA2. The three plots show a cell-level correlation analysis of the three markers for the entire gastruloid. Each dot in the correlation plot represents an individual cell. Shown are all three marker combinations. Quadrants show regions of background signal (-) and actual expression (+) for each marker in the respective color, defined by Otsu thresholding.

**Fig. 6.**
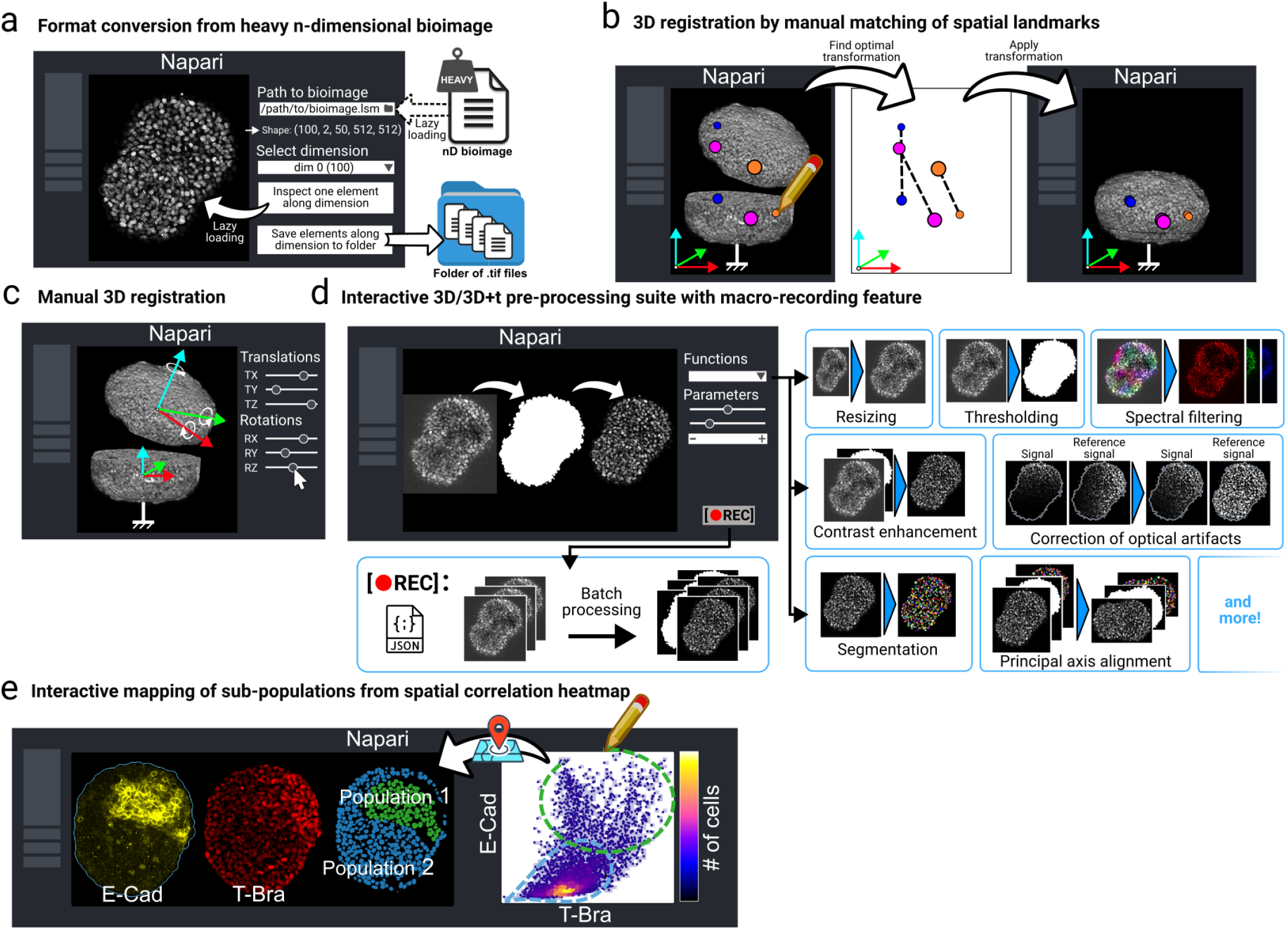
Interactive napari plugins for bioimage handling, manual registration, preprocessing, data exploration and analysis. **(a)** A napari plugin for exploring and processing large nD bioimages. Users can load datasets lazily, inspect slices along a chosen dimension, and export them as separate TIFF files. Compatible with common bioimage formats, it simplifies handling complex datasets. **(b**,**c)** To fuse data acquired with our dual-view setup, we provide a napari plugin to assist users in manually defining a 3D rigid transformation either when it is trivial (e.g a single axial translation) or to initialize an automatic registration tool close to the optimum. The first mode (left) allows the user to select salient landmarks and match them between the two views. An optimal rigid transformation is found automatically. The plugin’s second mode (right) lets the user define each element of the transformation (translations and rotations) until a satisfactory match is observed in 3D or on 2D planes. **(d)** A complete preprocessing suite is provided as a napari plugin to transform raw 3D and 3D+time datasets into datasets ready for analysis. The napari implementation allows for quick and visual exploration of parameters best suited to a given dataset. A recording feature can be toggled to save a complete user-defined pipeline into a *JSON* file, that can be used to process large datasets in batch or for sharing. **(e)** Interactive analysis of multiscale correlation heatmaps in napari is provided by allowing users to change the analysis length scale and to draw regions of interest directly on the correlation plot to see which cells contributed to the region’s statistics on a 3D view of the data.

When we applied the correction method to the T-Bra signal and evaluated re-normalized T-Bra intensity in a lateral section across the mid-plane of a 77 h gastruloid, we still observed a gradient with higher intensity in outer cell layers (Fig.5a), while the nuclei signal recovered a flat profile. Besides reflecting a true T-Bra expression gradient, a possible explanation for this observation could be hindered penetration of primary and secondary antibodies being considerably larger than Hoechst stain. To evaluate this, we compared immunostained T-Bra signal with the intensity of a T-Bra reporter (H2B-GFP expressed under a T-Bra promoter) in gastruloids grown from a transgenic cell line fixed at 72 h. We found a strong correlation between the two channels at all depths, excluding penetration artifacts (Fig.S7).

In addition to optical artifacts, our re-normalization method also corrects for intensity variations effectively induced by residual rotations of aggregates after flipping, which tilt the optical axis with respect to the axis of the first acquisition (Fig.2g). This is illustrated in Fig.5b, providing an example where the nuclei signal exhibits an asymmetrical gradient due to the fusion of two opposite views that were subject to residual rotations. After re-normalization, the nuclei profile is homogeneous, whereas the profile in T-Bra drastically changed, showing a polarization in the opposite direction with respect to the initial gradient that was also visible both in the nuclei stain and T-Bra channel. This emphasizes the importance of our method to properly quantify gene expression gradients in 3D. The observed polarized T-Bra appearance is in agreement with our previous study, in which we have reported that T-Bra expression already commences to be polarized at 77 h in some gastruloids [31].

Because the intensity decay in depth is wavelength-dependent (Fig.S3b), normalizing the different channels only based on the Hoechst intensity in the blue channel would have induced over-corrections in depth. We applied a wavelength-dependent correction to the different channels, by first measuring the characteristic length of exponential decay for the four colors independantly (Fig.S3a and c), and second by computing a map of Hoechst intensity corrected by this characteristic length. This gave us the equivalent of coarse-grained maps of nuclei intensity in each color, serving as a reference to correct the different channels (details in Methods III J).

We then applied the correction scheme to four-color 3D acquisitions of multiple gastruloids of different size (obtained by varying the initial cell number of the culture) imaged in the same slide, immunostained with T-Bra, the pluripotency marker Oct4, and the early endoderm marker FoxA2 (Fig.5c). We sorted these gastruloids into two groups and observed an approximately constant T-Bra expression for small aggregates (193 ± 20 *µm* average diameter, n=8), whereas big aggregates (291 ± 15 *µm* average diameter, n=5) showed a clear decay of T-Bra expression towards the interior (Fig.5c). Notably, we quantified T-Bra intensity as a function of distance to the gastruloid border using the *in toto* 3D image stack, thus reflecting expression gradients in 3D. For FoxA2, a similar pattern was observed, but expression appeared in much fewer cells (Fig.5d). Because FoxA2 signal was sparse and noisy, we could not compute simply the map of expression as done previously for T-Bra. Instead, we computed a map of the fraction of FoxA2 cells in a given neighborhood. We first thresholded the signal in segmented nuclei, using the interactive visualization tools developed in napari (see Fig.6 and Methods). This allowed us to identify the correct intensity threshold and generate a binary mask of FoxA2 expression. We then computed the number of FoxA2 positive cells rescaled by the local nuclei density, using the same method as applied for proliferation analysis, except that we consider FoxA2 positive cells instead of ph3 positive cells. The coarse-grained map of FoxA2 positive cells was then analyzed radially for gastruloids of various sizes, small gastruloids of 193 ± 20 *µm* average diameter (n=8) and bigger gastruloids of 278 ± 16 *µm* average diameter (n=3). We observed slightly higher FoxA2 levels in outermost cell layers, and in general more cells expressing FoxA2 in small gastruloids proportionally to the total number of cells. We applied the same method of radial analysis on 3D maps of Sox2 fraction, see Fig.S8a on 5 gastruloids at 60h. Cells expressing Sox2 are heterogeneously distributed but there is no radial gradient.Taken together, these analyses demonstrate that coarse-grained maps coupled with radial profiling can quantify large-scale patterns and gradients of gene expression across gastruloids.

Our ability to image multiple expression markers in the same acquisition without cross-talk and to reliably segment all cells allows correlating expression of different genes at the single cell level. To demonstrate this, we analyzed correlations between T-Bra, Oct4, and FoxA2 in the data described above, after spectral filtering and intensity renormalization. Both processing steps were crucial to obtain unbiased results (Fig.S8f,g). To discriminate marker positive from cells only showing background levels, we applied Otsu thresholding on the cells’ intensity distribution for each marker and confirmed the distinction by visible inspection.

We expected that FoxA2 and Oct4 are mutually exclusive (since cells are either in the process of endoderm differentiation or still pluripotent). A similar behavior was expected for T-Bra and Oct4/ T-Bra and FoxA2, although some co-expression could be present since T-Bra marks very early mesendoderm differentiation en route to meso- or endoderm and Oct4 is a relatively long-lived pluripotency marker [56].

We indeed observed that Oct4 and FoxA2 appeared as two mutually exclusive populations (Fig.5f, left). On the contrary, while the brightest Oct4 cells also showed low T-Bra signal and vice versa, a large pool of T-Bra and Oct4 positive cells appeared, not showing a striking correlation pattern for most cells (Fig.5f, center). For FoxA2 and T-Bra, a clear anti-correlation was observed, with cells appearing in all four quadrants (Fig.5f, right). Interestingly, we overall detected fewer FoxA2 positive cells than T-Bra positive cells. While we found many cells that were T-Bra positive and FoxA2 negative, most of the FoxA2 positive cells showed at least some T-Bra signal. This indicates that T-Bra expression might be required to activate FoxA2. However, cells expressing both markers still showed a negative correlation, suggesting that once FoxA2 is activated, T-Bra expression is repressed. Overall, these data suggest a sequence of cellular transitions from Oct4 progressively to T-Bra, from which a fraction of cells, presumably the ones expressing low T-Bra levels, will commit to FoxA2. To further explore such transitions and localize specific cell populations, tools that allow interactive mapping of cell populations selected in the correlation plot to the actual gastruloid are needed. We have, therefore, integrated this functionality into our napari plugin (see Fig6). We believe that our integrated imaging and analysis pipeline will allow researchers to study cell differentiation and cell state transitions at different stages of gastruloid development for the first time in full spatial context.

### Community-driven tool

We packaged our preprocessing and analysis pipeline into a *Python* library called *Tapenade* (for “Thorough Analysis PipEliNe for Advanced DEep imaging”). The code is freely available on *Github* at github.com/GuignardLab/tapenade/. The library comes with an extensive documentation and *Jupyter* notebooks accessible to non-specialist users. All datasets used to train our custom *StarDist* models to detect nuclei and mitotic events, along with the optimized weights for nuclei and mitotic events detection are currently available upon reasonable request. A non-specialist user can quickly segment a custom dataset by downloading the weights and loading them into one of the user-friendly implementations of *Stardist* like the *Stardist Fiji* plugin, the *napari* plugin *stardist-napari*, or the *ZeroCostDL4Mic* notebook [48, 57].

While working with large and dense 3D and 3D+time gastruloid datasets, we found that being able to visualize and interact with the data dynamically greatly helped processing it. During the preprocessing stage, dynamical exploration and interaction led to faster tuning of the parameters by allowing direct visual feedback, and gave key biophysical insight during the analysis stage. We thus created four user-friendly napari plugins designed around facilitating such interactions (Fig.6):

#### a. napari-file2folder

Efficient handling of large nD bioimaging datasets is a common challenge in bioimage analysis. To address this, we developed a napari plugin for lazy loading and efficient exploration of heavy bioimages using the zarr library. The plugin supports a wide range of standard bioimage file formats, including lsm, tif, ome.tiff, zarr, and czi. Upon lazy loading of an image, the user can immediately inspect its total shape across dimensions, even for datasets that cannot be fully loaded into memory. The plugin enables interactive selection of a single element along a specified dimension for focused exploration (e.g., viewing a single z-slice or a timepoint). Additionally, it offers a feature to save all elements along a selected dimension as separate tif files in a specified folder, which is particularly useful for downstream processing workflows with other plugins (Fig.6a). The plugin is freely available on the *napari* hub or at github.com/GuignardLab/napari-file2folder.

#### b. napari-manual-registration

When using our automatic registration tool to spatially register two views of the same organoid, we were sometimes faced with the issue that the tool would not converge to the true registration transformation. This happens when the initial position and orientation of the floating view are too far from their target values. We thus designed a napari plugin to quickly find a transformation that can be used to initialize our registration tool close to the optimal transformation. From two images loaded in napari representing two views of the same organoid, the plugin allows the user to either (i) annotate matching salient landmarks (e.g bright dead cells or lumen-like structures) in both the reference and floating views, from which an optimal rigid transformation can be found automatically using singular value decomposition [58] (Fig.6b), or (ii) manually define a rigid transformation by continually varying 3D rotations and translations while observing the results until a satisfying fit is found (Fig.6c). The plugin is freely available on the *napari* hub or at github.com/GuignardLab/napari-manual-registration.

#### c. napari-tapenade-processing

From a given set of raw images, segmented object instances, and object mask, the plugin allows the user to quickly run all preprocessing functions from our main pipeline with custom parameters while being able to see and interact with the result of each step (Fig.6d). For large datasets that are cumbersome to manipulate or cannot be loaded in napari, the plugin provides a macro recording feature: the users can experiment and design their own pipeline on a smaller subset of the dataset, then run it on the full dataset without having to load it in napari. The plugin is freely available on the *napari* hub or at github.com/GuignardLab/napari-tapenade-processing.

#### d. napari-spatial-correlation-plotter

This plugins allows the user to analyze the spatial correlations of two 3D fields loaded in napari (e.g. two fluorescent markers). The user can dynamically vary the analysis length scale, which corresponds to the standard deviation of the Gaussian kernel used for smoothing the 3D fields. If a layer of segmented nuclei instances is additionally specified, the histogram is constructed by binning values at the nuclei level (each point corresponds to an individual nucleus). Otherwise, individual voxel values are used. We took inspiration from the plugin *napari-clusters-plotter* [59] in letting the user dynamically interact with the correlation heatmap by manually selecting a region in the plot. The corresponding cells (or voxels) that contributed to the region’s statistics will be displayed in 3D on an independent napari layer for the user to interact with and gain biological insight (Fig.6e). The plugin is freely available on the *napari* hub or at github.com/GuignardLab/napari-spatial-correlation-plotter.

## II. DISCUSSION

### Multiscale imaging and analysis pipeline

We have developed an integrated pipeline for *in toto* imaging of whole-mount gastruloids and automated image processing. A key innovation of our approach is its specific optimization for dense 3D tissues, such as gastruloids, which present similar imaging challenges and artifacts as tumors and spheroids, unlike many organoids that are shallow and less light-diffusive. Unlocking deep whole-mount imaging is crucial for researchers working with such dense tissues, as cryosectioning strategies cannot fully substitute for comprehensive three-dimensional (3D) *in toto* analysis [18, 21]. Reconstructing 3D tissue microstructure from cryosectioned data is complex and challenging, involving slicing tissue into thin sections and imaging them individually. To achieve a 3D reconstruction, these 2D images must be meticulously aligned and stitched together, a process complicated by frequent issues such as tissue deformation and section loss. In contrast, whole-mount imaging preserves spatial relationships and tissue integrity, making it the method of choice for studying cellular interactions and tissue architecture in their native context. Additionally, whole-mount imaging reduces sampling bias and enhances the detection of rare cell populations or phenotypic variations that might be missed in thin tissue sections. This limitation also apply to state-of-the art spatial transcriptomic techniques, due to the fact that only few RNAs are typically captured per cell for each individual gene and many cells do not show any counts for a gene of interest expressed at low level [60].

### Application to other biological systems

In general, the pipeline is applicable to any tissue, but it is particularly useful for large and dense 3D samples, such as organoids, embryos, explants, spheroids, or tumors, that are typically composed of multiple cell layers and have a thickness greater than 50µm. The processing and analysis pipeline are compatible with any type of 3D imaging data (e.g. confocal, 2 photon, light-sheet, live or fixed). The applicability and limitations of the different modules of the pipeline are detailed Table I.

**TABLE I.** Purpose, requirements, applicability, and limitations of each step in the processing and analysis pipeline.

|  | Spectral filtering | Custom 3D nuclei segmentation | Normalization | Multi-scale analysis |
| --- | --- | --- | --- | --- |
| **Purpose** | Generate cross-talk free four-color images of markers imaged simultaneously | Quantifying cell-scale gene expression and nuclei morphological properties | Correct for intensity gradients due to optical artifacts in 3D | Analyze gene expression patterns, cellular events, and morphologies in 3D across multiple spatial scales |
| **Requirements** | Calibration of emission spectra as a function of imaging depth | Nuclei must be star-convex and $\sim 15$ px per nucleus after resampling | At least one ubiquitously expressed channel; calibration for wavelength-dependent correction | If domain-based signal (e.g. gene expression signals): domain expression mask. If object-based signal (e.g. morphological features) instance masks. |
| **Applicability** | Mostly useful in two-photon imaging because of broad excitation spectra | Tested on tissues from multiple species and modalities (Fig.S9d) | Applicable to any multichannel data across tissues and modalities | Any 3D data with gene expression or morphological information |
| **Limitations** | / | Small ($< 15px$/nucleus) and anisotropic resolution may impair segmentation | / | Hollow structures may bias results if no proper masking is available |

- Spectral unmixing to remove signal cross-talk of multiple fluorescent targets is typically more relevant in two-photon imaging due to the broader excitation spectra of fluorophores compared to single-photon imaging. In confocal or light-sheet microscopy, alternating excitation wavelengths often circumvents the need for unmixing. Spectral decomposition performs even better with true spectral detectors; however, these are usually not non-descanned detectors, which are more appropriate for deep tissue imaging. Our approach demonstrates that simultaneous cross-talk-free four-color two-photon imaging can be achieved in dense 3D specimen with four non-descanned detectors and co-excitation by just two laser lines. Depending on the dispersion in optically dense samples, depth-dependent apparent emission spectra need to be considered. Spectral filtering has already been applied in other systems (e.g. [61] and [45]), but is here extended to account for imaging depth-dependent apparent emission spectra of the different fluorophores. In our pipeline, we provide a code to run spectral filtering on multichannel images, integrated in Python. In order to apply the spectral filtering algorithm utilized here, spectral patterns of each fluorophore need to be calibrated as a function of imaging depth, which depend on the specific emission windows and detector settings of the microscope.
- Nuclei segmentation using our trained StarDist3D model is applicable to any system under two conditions: (1) the nuclei exhibit a star-convex shape, as required by the StarDist architecture, and (2) the image resolution is sufficient in XYZ to allow resampling. The exact sampling required is object- and system-dependent, but the goal is to achieve nearly isotropic objects with diameters of approximately 15 pixels while maintaining image quality. In practice, images containing objects that are natively close to or larger than 15 pixels in diameter should segment well after resampling. Conversely, images with objects that are significantly smaller along one or more dimensions will require careful inspection of the segmentation results. To evaluate our 3D nuclei segmentation model, we tested it on diverse systems, including gastruloids stained with the nuclear marker Draq5 from Moos et al. [62]; breast cancer spheroids; primary ductal adenocarcinoma organoids; human colon organoids and HCT116 monolayers from Ong et al. [29]; and zebrafish tissues imaged by confocal microscopy from Li et al [63]. These datasets were acquired using either light-sheet or confocal microscopy, with varying imaging parameters (e.g., objective lens, pixel size, staining method). Given the difficulty to access ground truth annotations of 3D organoid data and the multiplicity of models in the literature, we only show qualitatively the versatility of our segmentation model (Fig.S9). The results show satisfactory performance on these different datasets and imaging conditions.
- Normalization is broadly applicable to multicolor data when at least one channel is expected to be ubiquitously expressed within its domain. Wavelength-dependent correction requires experimental calibration using either an ubiquitous signal at each wavelength. Importantly, this calibration only needs to be performed once for a given set of experimental conditions (e.g., fluorophores, tissue type, mounting medium). To our knowledge, the calibration procedures for spectral-filtering and our image-normalization approach have not been performed previously in 3D samples, which is why validation on published datasets was not possible. Nevertheless, they are described in detail in the Methods section, and the code used from the calibration measurements to the corrected images is available open-source at the Zenodo link in the manuscript.
- Multi-scale analysis of gene expression and morphometrics is applicable to any 3D multicolor image. This includes both the 3D visualization tools (Napari plugins) and the various analytical plots (e.g., correlation plots, radial analysis). Multi-scale analysis can be performed even with imperfect segmentation, as long as segmentation errors tend to cancel out when averaged locally at the relevant spatial scale. However, systematic errors such as segmentation uncertainty along the Z-axis due to strong anisotropy may accumulate and introduce bias in downstream analyses. Caution is advised when analyzing hollow structures (e.g., lumen or curved epithelial monolayers with large cavities), as the pipeline was developed primarily for 3D bulk tissues, and appropriate masking of cavities would be needed.

In this work we have not used refined masks of the gastruloids that would account for and exclude hollow structures. In Fig.4(a) and (c), we noted that this lead to noticeable patterns in the packing-related coarse-grained fields and in the local alignment of the true strain tensor major axis. To improve our current implementation, our masking approach could be augmented by a second method used to specifically detect these structures. Pixel classification based on Random-Forests could be used to get accurate segmentation using only a few manually annotated lines. These approaches are now widely accessible to non-specialist users in softwares like *Fiji* (with *Labkit* [64]), *Ilastik* [65], or *napari* (with *napari-APOC* [66]).

### A Python integrated pipeline for non-specialists, with napari-enabled 3D exploration

We built our pipeline in Python to leverage its rich ecosystem of scientific computing libraries while keeping the code as accessible and user-friendly as possible. We provide detailed installation instructions and Jupyter notebooks with step-by-step instructions designed to be used by non-specialists, with a focus on ease of use and reproducibility. To tackle the challenges that arise from manipulating deep and dense 3D data, we provide napari plugins for manual registration, preprocessing, and analysis. napari provides a user-friendly graphical interface which enables real-time visualization and exploration of 3D data during each step of the preprocessing and analysis. This was essential for the development of the pipeline, as the direct visual feedback allowed us to quickly iterate and optimize the processing steps, while gaining insights into the 3D organization of gastruloids. We believe our pipeline offers a distinctive and effective integration of 3D data exploration and processing, providing a valuable tool for the organoid and tumoroids research community.

We did not explore continuously adjusting laser power based on excitation depth to enhance signals in deeper tissue layers, which could further extend the depth limits of our imaging method. In principle, incorporating such adaptation should not affect our pipeline, as long as the laser used for exciting the nuclei channel (which normalizes all other signals) is the same or maintains the same power-to-depth relationship as the lasers exciting the other fluorophores to be normalized. Here, we developed a wavelength-dependent normalization scheme based on a simplified columnar model. Future work could focus on creating a complete model that accounts for photon paths in a complex geometry, for example in elongated gastruloids, and integrate the spectral filtering approach to the wavelength-dependent normalization scheme.

In this work, we assume that the nuclear signal is the most reliable for normalizing other signals due to its extensive coverage of the tissue (over 60% in gastruloids). Instead of normalizing signals at the level of individual nuclei, we developed a correction scheme that identifies the optimal scale of characteristic gradients in nuclear intensities to generate a continuous field for re-normalization. This approach has the added advantage of being applicable to markers with different subcellular localizations (e.g., nuclei, cell membrane, intracellular organelles).

A necessary component of our quantitative imaging pipeline was the use of spectral unmixing to address spectral cross-talk, an inherent challenge in multi-color acquisitions, particularly with two-photon excitation. This allowed us to image four different fluorescent targets simultaneously and was required to capture and correlate multiple genetic markers on the single-cell level. This can be further expanded in the future by using state-of-the-art hyperspectral detection, which allows capturing up to 32 channels simultaneously and thus decompose even more spectrally overlapping fluorescent targets [67]. Two-photon excitation is advantageous in this context, allowing for the excitation of multiple targets with a single laser line. In our work, three of the four fluorophore species (Hoechst, AF488, and AF568) were excited using a 920 nm laser line. Additionally, iterative rounds of immunostaining and washing can be applied to maximize the information content [24, 68]. In this context, the interactive registration tools that we have implemented here will be key ingredients to correct for sample movements between subsequent staining steps.

### Exploring gastruloid development in 3D

In the context of gastruloid development, which serves as the central biological theme of this paper, it is crucial to understand how spatial heterogeneities in gene expression arise at both the cell microenvironment scale and the larger gastruloid scale. During early gastruloid symmetry breaking and elongation, radial and later antero-posterior gradients of the gene T-Bra, a nascent mesoderm marker, emerge. These radial gradients can be captured with our coarse-grained analysis (see Fig.5c). However, while both the endoderm marker FoxA2 and T-Bra exhibit radial gradients at the coarse-grained scale, their anti-correlated expression is only evident at the single-cell level (see Fig.5c-f). This example highlights the importance of a multiscale analysis pipeline that allows simultaneous visualization of tissue-scale properties and local cellular events. Our methodology utilizes segmentation of the Hoechst-stained nuclei channel to (1) identify individual cells for cell-scale analysis and (2) normalize other channels using the nuclei signal. By developing a diverse training set for nuclei segmentation, we achieved precise determination of nuclei shapes. This precision enables systematic studies of the relationship between nuclear morphology and gene expression patterns, and can be extended to cell morphology using membrane stains such as ZO-1 and cell shape segmentation methods like CellPose [69, 70]. Note that the latest version of the latter method, CellPoseSAM [70], can give satisfactory results for nuclei segmentation, and we showed qualitatively that it benefits from the Tapenade preprocessing workflow for segmenting deep tissues (Fig.S10a) and that using it as a replacement for our trained StarDist3D model lead to similar results for the nuclear morphology study (Fig.S10b-d).

Nuclei deformation patterns show large-scale alignment at the gastruloid level (see Fig.4c-d) and localized alignment near lumens and cavities, providing potential insights into the structuring of epithelial and mesenchymal tissues during gastruloid development. In older gastruloids with cavity-containing structures, such as somitic structures [38], automated cavity detection will be crucial for accurately quantifying coarse-grained fields of cell or division densities. While this work focuses on gastruloid datasets, we believe that the pipeline effectively extends to organoid, tumoroid, and other datasets that encounter challenges related to whole-mount deep imaging. We designed the library in the prospect of long-term maintenance, and it will receive continuous improvements in the future.

## III. MATERIALS AND METHODS

### A. Sample preparation

#### 1. Gastruloid culture

Gastruloids were generated using the protocol described previously in [31], from four different cell lines. Briefly, cells were seeded and aggregated for 48 h in low-adherence 96-well plates (Costar ref: 7007) and subsequently pulsed with the Wnt agonist Chiron, which was washed out after 24 h, i.e. at 72 h of aggregate culture. Time points indicated throughout the paper refer to the time after seeding. The two cell lines T-Bra-GFP/NE-mKate2 and E-cad-GFP/Oct4-mCherry were used for gene expression analyses. The E14Tg2a.4 line (ATCC) was used for spectral calibration and benchmarking as it is devoid of any fluorescent construct. The H2B-GFP (a generous gift from Kat Hadjantonakis) was used for live imaging experiments and morphometric analyses. In brief, between 50 and 400 cells were seeded in low adhesive 96 wells plates in a neural differentiation medium. In order to promote endoderm formation and to reduce symmetry breaking variability compared to the standard gastruloids protocol [55], we enriched the medium with Fibroblast Growth Factor and Activin as in [31]. All cell lines were tested to be free from mycoplasma contamination using qPCR.

#### 2. Immunostaining

Samples were immunostained using the same protocol as in [31] apart from the fact that samples were incubated in a 1 mM glycin solution, and that primary antibodies (see table below) were incubated during 3 consecutive days. Secondary antibodies AF488, A568, AF647 (ThermoFisher) and Hoechst 33342 (ThermoFisher) were chosen to be compatible with four-color two-photon imaging. They were diluted 500 times for incubation, except of Hoechst which was diluted 10000 times in order to minimize the crosstalk of the Hoechst signal into other channels.

| Antibodies |  |  |  |  |
| --- | --- | --- | --- | --- |
| Antibody | Species | Reference | Provider | Dilution |
| Brachyury | Goat | AF2085 | RD system | 1:40 |
| E-cadherin | Rat | M108 | Takara | 1:200 |
| FoxA2 | Goat | sc-6554 | Santa-Cruz | 1:50 |
| Snail | Goat | AF3639 | RD system | 1:50 |
| OB-cadherin | Rabbit | 71-7600 | Invitrogen | 1:500 |
| pH3 | Rat | ab10543 | abcam | 1:250 |
| Oct4 | Mouse | Merck<br>Millipore | MAB4419 | 1:100 |
| Sox2 | Mouse | SantaCruz<br>Biotechnology | sc-365823 | 1:100 |
| anti-GFP | Chicken | Aves | GFP-1020 | 1:100 |

#### 3. Mounting

Samples were mounted between two glass coverslips of standard thickness (0.13 to 0.17 mm) using 250 *µm* or 500 *µm* spacers (SUNJin Lab, Taiwan, R.O.C.) in 20 *µl* or 40 *µl* mounting medium. For the clearing benchmark, gastruloids were incubated overnight and then mounted in either PBS, 80% glycerol solution in PBS (v/v) [41], 20% optiprep solution in PBS (v/v) [42], or antifade mounting medium (ProLong Gold Antifade Mountant, Thermo Fisher Scientific). For all other experiments with fixed samples, gastruloids were mounted in 80% glycerol solution in PBS (v/v). For live imaging, gastruloids were transferred from 96-well plates to either MatTek dishes (MatTek corporation, ref: P35G-1.5-14 C) or to micro-well plates (500 *µm* wells, SUNBIOSCIENCE ref: Gri3D) in culture medium and imaged in a chamber maintained at 37 ^*°*^C, 5% *CO*2 with a humidifier.

### B. Two-photon imaging

Four-colors two-photon imaging of immunostained samples was performed on an upright Nikon A1 R MP microscope using an Apo LWD 40 × /1.1 WI *λ*S DIC N2 objective. Fluorescence was simultaneously excited at 920 nm (to excite simultaneously Hoechst, AF488 and AF568) with a tunable femtoseconde laser (Chameleon Discovery NX from Coherent) and at 1040 nm (to excite simultaneously AF568 and AF647) with the same laser secondary output at fixed wavelength. Fluorescence was detected on four non-descanned GaAsP detectors with the following emission filters: 450/70 (Ch1), 530/55 (Ch2), 607/70 (Ch3), 700/50 (Ch4). To spectrally separate the channels, three dichroic mirrors (488 LP, 562 LP, 650 LP) were used. Multiposition imaging was used to automatically acquire image stacks on multiple gastruloids mounted in the same sample slide.

Live imaging of gastruloids was performed on a Zeiss 510 NLO (Inverse - LSM) with a femtosecond laser (Laser Mai Tai DeepSee HP) with a 40 ×/1.2 C Apochromat objective. The Histone-GFP signal was excited at 920 nm and detected with a non-descanned GaAsP detector with a 560 LP. The images were acquired with the full field-of-view (318 *µm*, pixel size 0.62 *µm*), and a z-spacing of 1 *µm*. Depth in z varies from 150 to 350 *µm*, depending on the acquisitions and the sample size, but always past the midplane of the gastruloid.

### C. Confocal imaging

Confocal imaging was performed on a Zeiss LSM880 system (Carl Zeiss) using a Plan-Apochromat 40 ×/1.2 NA water immersion objective. Fluorescence was excited with the 488 nm line of an Argon laser. To split excitation and emission light, a 488 dichroic mirror was used. Fluorescence was detected between 491 and 700 nm on a 32-channel GaAsP array detector, with a pinhole set to one airy unit.

### D. Image quality estimation

To quantitatively assess clearing efficiency and compare the information content of images at depth, we used the Fourier ring correlation quality estimate (FRC-QE) [43]. We selected a column of 400 ×400 pixels for the clearing benchmark and 100 × 100 pixels for all other analyses in the center of aggregates and computed FRC-QE using the available Fiji plug-in.

### E. Preprocessing: Spectral unmixing

To remove spectral cross-talk, four-channel images were spectrally unmixed into *n* images corresponding to the contribution from each of the *n* fluorophore species present in the sample.

#### 1. Spectral calibration

To this aim, reference spectra 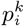of the utilized fluorophores were determined from calibration images acquired on aggregates stained with only one fluorophore species at the same excitation and detection settings. To account for wavelength dependent signal decay with imaging depths, spectral patterns were computed as function of imaging depth. It is important to note that spectral patterns depend on the gain settings of each detector. However, once calibrated for a certain set of gain values, reference patterns for different gains can be easily interpolated by simply measuring the count ratios for both sets of gains on a sample providing sufficient signal (i.e. well above background counts) on each detector for all gains.

#### 2. Spectral decomposition

Four-channel images *I*^*k*^(*x, y, z*) were decomposed using a previously published unmixing algorithm originally developed in the context of fluorescence lifetime correlation spectroscopy [71], image correlation [72] and fluorescence correlation spectroscopy [45, 73]. First, spectral filter functions 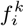 were calculated using the calibrated reference spectra 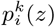:

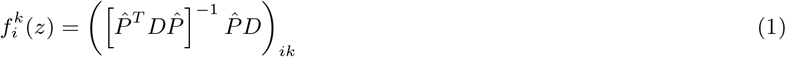

Here, *P* is a matrix with elements 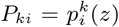 and *D* a diagonal matrix computed from the average image intensity in each channel *k, D* = *diag* [1*/*⟨*I* ^*k*^ (*x, y, z*)⟩*x*,_*y*_].

The spectral filter functions represent intensity weights per spectral channel. To obtain the filtered images *Ii*(*x, y, z*) for each fluorophore species *i*, the channel images are multiplied with the filter functions and summed over all channels (*m* = 4 here):

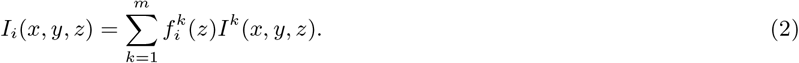

It should be noted that unmixing successfully works as long as signal levels are well above background counts and all images, acquired for spectral calibration and in actual imaging experiments, are devoid of detector saturation.

This way, even closely overlapping fluorophore species can be discriminated with only few detector channels, at the expense of increased image noise [72]. In this context, pure detector channels, i.e. channels that predominantly detect one of the fluorophore species, are beneficial. This is the case in the example of GFP and AF488 presented in Fig.S1e,f, in which the first channel detects mostly GFP fluorescence. In addition, samples in which signal from different fluorophore species is detected in different pixels (e.g. pixels corresponding to nuclei and membranes) are generally easier to unmix and suffer less from noise increase than samples were different fluorophore species contribute to the signal in the same pixels.

### F. Preprocessing: Dual-view registration and fusion

Multiple organoids were acquired from the same slide, first from one side for all of them, then from the other side. Therefore, the first step of the registration and fusion algorithm was to pair the two sides of each organoid together. This pairing was done by extracting the sample positions stored in the metadata of the acquisitions and finding the pairing that minimizes the sum of the distances between paired samples using a linear assignment algorithm.

Registration of these dual-view image stacks was achieved using an existing intensity based blockmatching algorithm [44], adapted for 3D microscopy images and previously used in [6, 7]. From the two distinct images, one is arbitrarily chosen as the reference image and the other, named the floating image, is registered onto the reference one. Starting from an initial transformation, the blockmatching algorithm compares, on multiple hierarchical levels, each block in the floating image with the corresponding neighboring blocks in the reference image, and finds the rigid 3D transformation (specified by rotation around axes X, Y, Z and translation along axes X, Y, Z) from these sets of blocks. Then, we apply the transformation to the floating image, so that the two views are in the same referential, and fuse them into one image stack.

Because of the computational cost of comparing blocks in large 3D images, the size of the neighborhood that the registration algorithm explores is restricted. Therefore, the magnitude of the transformation between the two image stacks has to be relatively small (typically less than 30^*°*^ rotations and 100 units in translation). As a consequence, the user has to provide an initial transformation to the algorithm, which approximates the actual transformation. In the case of dual-view imaging with opposite views, the initial transformation would correspond to 180^*°*^ rotation around the X axis and some additional translation. To explore the initial transformation, we developed the napari plugin *napari-manual-registration* detailed above and illustrated in Fig.6. This tools allows to interactively rotate and translate the floating with respect to the reference image stack. The tool also allows users to specify manual landmarks in both image stacks, e.g. lumen, dead or dividing cells, from which an initial transformation can be automatically computed.

While the blockmatching algorithm generally allows for any type of transformation (rigid, affine, non-linear, …), we restrained the search space to rigid transformations. Moreover, since the stacks comprise multiple channels, we used a reference one that was expressed ubiquitously, here the nuclei staining, on which the transformation was computed. The other channels were registered using the same transformation matrix as the reference channel. Because the SNR of each stack decreased roughly with the distance to the objective, we weighted the contribution in the fusion of the two opposing sides of the sample. The weights *f*1 and *f*2 respectively for the reference view and floating view were computed as a sigmoid:

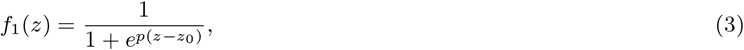

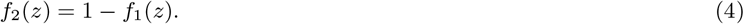

with *z* being the depth in the stack in voxels, *z*0 the middle (i.e inflection point) of the sigmoid, where *f*1(*z*0) = 0.5, and *p* the slope of the sigmoid. The decay length *l* of the sigmoid and the effective fusion width δ are linked by *l* = 4*/p* and δ = 2*l*.

As illustrated in Fig.2e, the two sides were imaged up to a certain depth depending on the sample size and the image quality, past the mid-plane to ensure some overlap with the other view. Because we acquired the same depth and number of z-planes for both views, the sigmoid is centered in the middle of the overlapping region, which is why we considered *z*0 = 0.5. The overlapping region to fuse had a typical width value of *w* = 70 to 150 *µ*m (depending on the image depth) around the midplane. The coordinate along the overlap region was mapped to the range [0, 1] to allow setting the parameters of the sigmoid fusion function independently from the size of the overlap or the size of the gastruloid, as illustrated on Fig.S5. The slope of the sigmoid was set to *p* = 15, which leads to *l* = 0.27 and δ = 0.53. Intuitively, this means that 53% of the overlapping region was used to mix both signals with non-negligible mixing coefficients to create the fused part.

### G. Nuclei segmentation

To segment nuclei, we trained a custom Stardist3D [48] model on three annotated datasets acquired with two-photon microscopy in different acquisition modalities: one fixed dateset, two live datasets (one with mosaic fluorescence labeling), see Table II and S2a.

**TABLE II.** Description of the datasets used for training our custom StarDist3D model to detect nuclei. All datasets were acquired with a two-photon microscope.

| density | dimension | num. nuclei | pixel size ($\mu\text{m}/\text{pix}$) |
| --- | --- | --- | --- |
| dense | 3D | 1018 | Z: 1, Y/X: 0.25 |
| dense | 3D+time | 100 | Z: 1, Y/X: 0.62 |
| sparse | 3D+time | 3296 | Z: 1, Y/X: 0.62 |

#### 1. Global contrast enhancement

Before training and inference with our Stardist3D model, we applied percentile-based histogram clipping and normalization to the input images. For a given image voxel at position 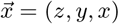, its intensity 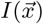 is first transformed as follows:

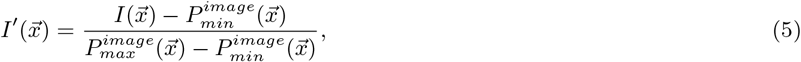

where 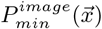 and 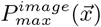 are low and high percentile intensity values in the image.

In practice, we choose 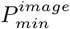 and 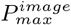to be the first and 99th percentile of the intensity values in the image. As shown Fig.S2d, applying global contrast enhancement was not sufficient to restore sufficient intensity in the deeper regions of some samples, which lead to poor segmentation performances. We thus extended our contrast enhancement method to apply in local sub-regions of the images.

#### 2. Local contrast enhancement

Because the image contrast is not homogeneous within and across samples, we homogenized intensity profiles across the training datasets by applying local percentile-based histogram clipping and normalization.

For a given image voxel at position 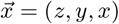, its intensity 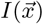 is first transformed as follows:

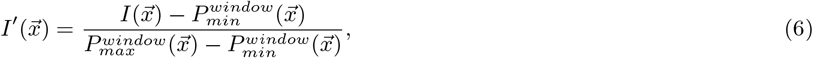

where 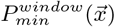and 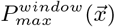 as before, the first and 99th percentile of the intensity values within the window, and *w* close to the average size of the nuclei in the dataset The transformed image *I*^*′*^ is finally clipped to the range [0, 1] to obtain the locally enhanced image.

The training datasets were rescaled to an isotropic voxel scale of 0.62 µ*m/pix* in all directions, from which crops of 64 × 64 × 64 voxels were extracted and augmented during training by 3D rotations, flipping, intensity shifts, and additive Gaussian noise. Applying scaling transformations to crops during training is often regarded as the best way to achieve robustness in the case of a wide object size distribution. This is typically the case when several raw datasets acquired in different experimental conditions are combined, particularly due to different pixel sizes across datasets. However, we found in our case that the size distribution related to a single dataset is narrow enough that robustness is better achieved by first rescaling all training datasets to a common isotropic nuclei size. [69, 74].

Before inference, a new dataset simply needs to be rescaled to reach that common nuclei size.

### H. Mitotic cells segmentation

Another StarDist model was trained to detect the divisions stained by ph3. To this aim, we iteratively trained StarDist3D models with a very small batch of annotated data, using the model trained on all nuclei, and corrected the prediction using napari to train again. At the end, 31 images of whole-mount gastruloids and 5300 divisions were used to train a custom Stardist3D model, using the same parameters as for the nuclei detection model in terms of preprocessing, crops size and data augmentation.

Whereas the model trained on nuclei was not very successful on ph3 stained nuclei, because of their particular shapes, the final model after multiple iterations achieved better results, around F1=82 %. Some over-segmentations around large and deformed nuclei were still present, and using 3D Gaussian blur did not achieved a better score, because they led to misdetections where neighboring nuclei were in mitosis. Therefore, after prediction, post-processing was used to filter out small volumes with thresholding. Events detected from the background or dead cells (using a mask of the sample computed from the nuclei channel) and minimal manual curation was done, using napari.

### I. Masked Gaussian convolution to probe spatial fields at different scales

To probe volumetric signals at different length scales *σ* ranging from the average nuclei size to the width of the organoid, we convolve signals with a Gaussian kernel of size *σ*. We propose two ways to compute convolutions depending on whether the signal is dense (e.g an image representing a signal intensity) or sparse (e.g nuclei volumes defined only locally on nuclei centroids).

#### a. Dense data

We apply convolution with a normalized Gaussian kernel to dense signals. For a given voxel at position 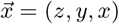, its value 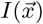, whether scalar or vector, is transformed according to:

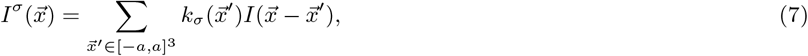

with 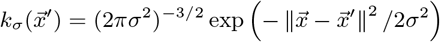, and *a* the convolution window size. In our implementation, we choose *a* = 3*σ*.

When a binary mask (e.g defining the inside of the organoid) is available, we apply a masked convolution with a correcting normalization factor. Let *M* be a binary masking function, e.g 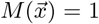 for voxels at positions 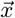 inside the organoid, and 0 everywhere else, we compute

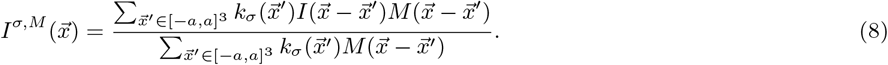

This additional normalization factor prevents the artifactual decay of the convolved signal close to the mask boundaries by “excluding” voxels outside the mask from the convolution.

Note that the masked formulation allows the user to compute values of a quantity outside its domain of expression. For instance, if both a mask *M*_*inst*_ of nuclear instances and a mask *M*_*in/out*_ that marks the inside and outside of the sample are provided, a quantity like the nuclear Hoechst expression can be computed in the full region defined by *M*_*in/out*_, even though it only appears in the nuclei.

#### b. Sparse data

For sparse data, i.e quantities defined locally at *n* specific positions 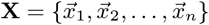, we compute the convolution and masked convolution in a similar manner, but with a mask *M* that is non-zero only at the positions at which the sparse signal is defined (e.g nuclei centroids). This formulation allows the user to choose between studying the sparse signal (i) at its original positions, i.e by computing the above convolution at positions 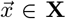, (ii) at resampled positions, e.g on a uniform grid for convenience, or (iii) as a dense continuous field across the whole domain of the gastruloid.

We provide a second implementation of the masked convolution specifically for sparse data, which is more convenient when the smoothed sparse signals should also be smooth (e.g when it should be defined at its initial positions or on a regular grid). This implementation uses the Kd-Tree data structure implemented in *Scipy* to efficiently find neighbor relationships from the positions at which the sparse signal is defined, which is used to optimize the computation of the convolution (8).

### J. Preprocessing: Intensity normalization

Quantifying gene expression in 3D tissue samples from imaging data is hindered by various sources of noise and optical artifacts that induce large scale intensity gradients (e.g. scattering, aberrations). The main sources of artifacts are (local) variations in optical paths and cell density, that usually manifest as decreased intensities towards deep tissue regions. Artifactual gradients can also appear during the registration of two non-collinear views, which further complexifies the expected distribution of artifacts and resulting intensity gradients.

We model the intensity signal 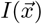 from a given channel as

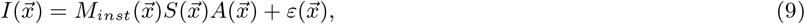

where 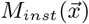 is a binary mask of the signal instances (e.g nuclei or membranes), i.e 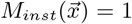 if 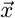 is inside an instance and 0 otherwise, 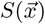 is the gene expression field, which provides a continuous representation of the cell-scale variations of the biological signal, 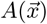 is the field representing the artifactual intensity variations, and 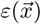 is an additive uncorrelated noise term which models both variations in the signal at scales below the cell-scale and acquisition noise. This decomposition is illustrated on Fig.S4.

We hypothesize that the Hoechst signal 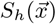 is globally homogeneous throughout the volume, i.e 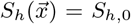, and that the field of optical artifacts *A* has large scale variations (typically superior to a few cell diameters). In particular, direct observations of homogeneous signals like Hoechst with different emission wavelengths, as displayed of FigS3(a-b), indicate that optical artifacts are mostly dominated by exponential decay with depth with a wavelength-dependent decay length. A second order contribution comes from scattering effects related to heterogeneities in photon’s optical paths, cellular density, or to geometrical considerations, e.g proximity to the sample’s boundaries. For every wavelength *λ* and every signal, we thus model *A* to first order as

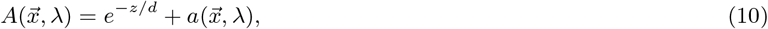

with *d* the wavelength-dependent decay length. The second term *a* on the right hand side corresponds to second order corrections to the exponential decay model. Note that, in accordance with Beer-Lambert law, the value *d*(*λ*) also likely depends on experimental parameters ℰ, like the nature of the mounting medium or the settings of the experiments, in a multiplicative fashion, i.e

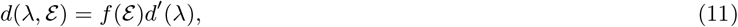

so that in similar imaging scenarios, ratios of decay lengths at two wavelengths

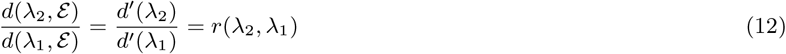

only depend on *λ*1 and *λ*2.

To estimate the decay lengths across the range of wavelengths in use, we measured the intensity decay in samples with ubiquitous nuclei stainings in four emission wavelengths: Hoechst for the blue channel (*λ* = 405*nm*), H2B-GFP amplified for the green channel (*λ* = 488*nm*), SPY-DNA555 or the endogenous H2B-Tomato for the red channel (*λ* = 555*nm*) and DRAQ5 for the far-red channel (*λ* = 647*nm*). Stainings for different wavelengths were done on separated samples to avoid signal crosstalk between the channels. All samples were imaged with the same excitation and detection settings. For each sample, we extracted the intensity profile in a central column along the z-axis to minimize boundary artifacts, and fitted equation (10) to recover the decay lengths *d*(*λ*) (Fig.S3(a-c)). We observe a monotonous relationship between median values of *d* across different samples and *λ*. To confirm our measurements, we also computed decay lengths via a second method: notice that the model of equation (10) leads to the following identity for two ubiquitous signals *I*1 and *I*2 at wavelengths *λ*1 and *λ*2 (neglecting the noise term *ε* in equation (9) and the second order term from equation (10))

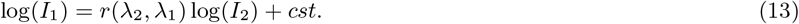

We imaged new samples with both Hoechst (*λ* = 405*nm*) and the Far-red nuclei stain (*λ* = 647*nm*). There was no significant spectral overlap between the two signals. By fitting equation (13), we obtained another estimate of the ratio *r*(*λ* = 405*nm, λ* = 647*nm*) ≈ 0.46, consistent with the value of 0.48 obtained with the previous method (Fig.S3e). In Fig.S3(d), we plot the decay length ratios with the decay length of the Hoechst channel *d*_*blue*_. In our model, these ratios are independent of imaging conditions, and the monotonicity and smoothness of the trend suggests that the ratios *d*_*blue*_*/d*(*λ*) in the range *λ* ∈ [405*nm*, 647*nm*] could be estimated by interpolating our results.

In our experiments, it was not possible to image ubiquitous nuclei signals at the same wavelengths as relevant biological signals, while Hoechst staining was often easily available. We thus developed a correction scheme based on Hoechst staining to separate the contribution of artifactual gradients from true gradients in gene expression. The rationale of the scheme stems from the fact that equation (13) also allows us to relate two ubiquitous signals up to a multiplicative constant:

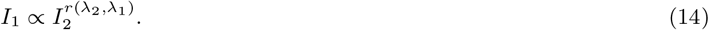

The Hoechst signal could therefore be used to predict the intensity profile from another “virtual” ubiquitous signal imaged at some wavelength *λ*≠*λ*_*blue*_ that coincides with the wavelength of the biological signal of interest.

First, we compute the Gaussian average of the reference Hoechst signal *I*_*h*_ at a scale *σ* using equation (8), in which we keep only voxel intensities that belong to the nuclei using a mask *Mnuc*:

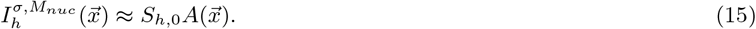

Here, *σ* is chosen large enough to average out the additive noise *ε* but small enough that the structure of the field of optical artifacts *A* is preserved. The practical choice of *σ* is discussed below.

We then normalize other biologically relevant intensity signals *I*_*b*_ by the Gaussian-averaged nuclei field 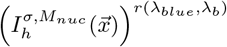 to get

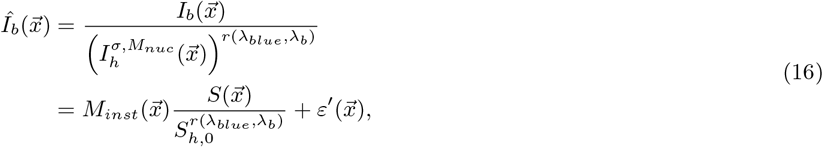

with *ε*^*′*^ the normalized version of the additive noise *ε*. This step effectively removes the field of optical artifacts from the signal. The choice of *σ* used for the Gaussian averaging of the ubiquitous signal depends on the length scale at which the field of artifacts *A* varies. Since this is not known in practice, we choose in our implementation the value *σ* that leads to the most homogeneous normalized ubiquitous signal across all z-planes. Formally,

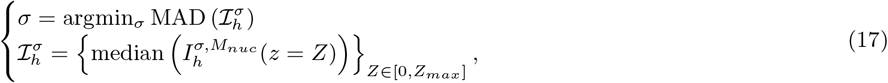

with MAD the median absolute deviation and *Zmax* the depth of the last z-slice. Fig S3f shows the decay profile of a sample normalized using different values of *σ*, as well as the corresponding Gaussian-averaged nuclei fields.

With the aim of comparing normalized signals across different samples acquired with the same imaging setup, we further rescale the normalized signal by a global multiplicative factor designed to mitigate the sample-dependent impact of the normalizing factor 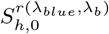 in equation (16). We multiply each normalized signal by a multiplicative factor such that the median intensity of the raw Hoechst signal computed in a region of brightest intensity is transmitted to the normalized Hoechst signal.

The rationale is that the median intensity in the brightest regions reflects the expected intensity in the absence of artifacts, e.g. in outer cell layers close to the cover glass.

We tested the normalization method on a sample stained with both Hoechst and a Far-red nuclei stain, using Hoechst to normalize the Far-red channel and then look at the intensity profile. Fig.S3e shows the depth decay for the unnormalized signal, as well as the signal normalized using the ratio *r*(*λ*_*blue*_, *λ*_*far*−*red*_) = 0.48 as found experimentally. On the same plot we show the profile for the ratio *r* = 1 that would model a situation where we do not correct for wavelength dependency. In the latter, the intensity is over-corrected in depth, whereas with the correct ratio we recover a flat profile in depth. Because there is an error associated with the measurement of *r*, we tested small variations around the value 0.48. The depth decay changes slightly, as shown on the right panel with a 10% difference above and below. XZ profiles for these two panels are shown on the right. In addition, we correct the Hoechst for its own intensity decay, in that case using *r* = 1 which recovers a flat profile in depth, this is shown Fig.S3g with the corresponding XZ views. The last sanity check is performed by plotting the profile not only along the Z axis but also along X and Y, to ensure the absence of any other gradients of intensity. This is Fig.S3h, plotted on the Far-red channel normalized with *r* = 0.48.

### K. Density maps

To quantify the local abundance of a target object, e.g. nuclei or specifically mitotic nuclei, we study three spatial fields: the density (number of objects per volume), the volume fraction (percentage of local space filled by the objects), and the map of objects volumes.

All fields are first computed at the voxel scale. The voxel-scale density field and map of objects volumes are computed by replacing each object by a single voxel of value, respectively 1 or the object’s volume placed at its centroid. The voxel-scale volume fraction corresponds to a binary mask of the target objects, and can be obtained directly with semantic segmentation methods or by binarising segmented object instances as output by Stardist3D. The voxel-scale fields are finally convolved with a Gaussian kernel of the desired analysis scale.

These quantities give a complete description of the local object abundance, as the volume fraction and map of objects volumes inform on the impact of volume on the object density. For instance, a large spatial region can misleadingly display a constant volume fraction but have large variations in object density and local objects volumes if the latter two compensate.

### L. Morphometric analysis

To quantify object-scale deformation, we fit an ellipse to each segmented object. This can be done efficiently by computing the inertia tensor *I*. For a given object composed of *N* voxels at positions 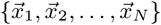, we first compute the positions of the voxels relative to the object’s centroid 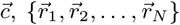, with 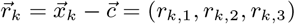. The inertia tensor is then defined as:

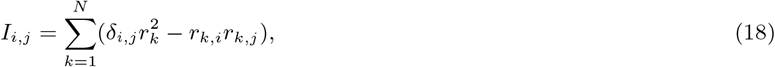

with 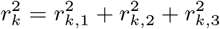 and δ*i*,*j* the Kronecker delta. The eigenvectors and eigenvalues of the inertia tensor correspond to the principal axes and moments of inertia of the object. Notably, the principal axes of *I* are the same as the principal axes of the ellipse that best fits the object. The lengths of the ellipse’s semi-axes can be obtained from the moments of inertia: if (*I*1, *I*2, *I*3) are the moments of inertia associated with the principal axes 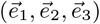, the lengths of the corresponding semi-axii (*L*1, *L*2, *L*3) are given by:

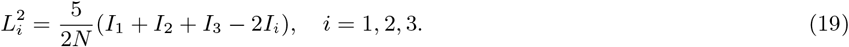

Computing the inertia tensor for each object leads to a sparse tensor field that can be studied at arbitrary scales using the sparse Gaussian convolution equation defined previously. For instance, it can be used to study local object alignments by focusing only on the largest semi-axis (the main axis of deformation of the object). In practice, quantifying local object alignment by averaging inertia tensors introduces a bias by over-representing large objects. To avoid this bias, we instead compute the true strain tensor from the inertia tensor. The true strain tensor *T* is defined as:

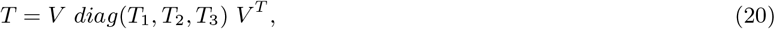

with *V* the matrix of eigenvectors of *I* and 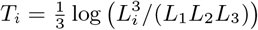. The eigenvalues (*T*1, *T*2, *T*3) of *T* represent the relative amount of deviation of each principal length from the radius of a sphere with the same volume as the object. Due to its relative nature, the true strain tensor is suited to compare the deformations of two objects with different sizes, and is thus appropriate for averaging in a local neighborhood.

## IV. CODE AND DATA AVAILABLITY

All datasets used to reproduce the figures of the articles, the trained Stardist3D weights, and the associated training datasets can be found in the following link: https://zenodo.org/records/14748083

The Tapenade package is available on GitHub: https://github.com/GuignardLab/tapenade.

The napari plugin napari-manual registration is available here: https://github.com/GuignardLab/napari-manual-registration

The napari plugin napari-tapenade-processing is available here: https://github.com/GuignardLab/napari-tapenade-processing

The napari plugin napari-spatial-correlation-plotter is available here: https://github.com/GuignardLab/napari-spatial-correlation-plotter

## ACKNOWLEDGMENTS

This work is supported by the French National Research Agency (“France 2030”, ANR-16-CONV-0001 from Excellence Initiative of Aix-Marseille University - A*MIDEX), a generic grant to P.-F.L. (ANR-19-CE13-0022), a generic grant to S.T. ANR-22-CE30-0021 and an ERC grant (to P.-F.L., ERC SyG 101072123). We also acknowledge the France-Bioimaging Infrastructure (ANR-10-INBS-04). This work was supported by the Fondation pour la Recherche Médicale, (to A.G, grant number FDT202404018538, and to P.-F.L., grant number EQU202003010407) V.DE. acknowledges support by an HFSP long-term postdoctoral fellowship (HFSP LT0058/2022-L). We thank Dalia El-Arawi for providing one of the datasets used in Fig.S10.

## CONTRIBUTIONS

S.T, P-F.L and L.G designed the project. S.T and V.D-E designed the experiments. A.G, V.D-E, S.T and A.R conducted the experiments and collected the data. V.D-E developed the spectral filtering approach, A.G and J.V annotated nuclei data for the training of the StarDist3D network. A.G and L.G adapted the registration algorithm, with input from J.V. and V.D-E. J.V developed and packaged the pipeline and the napari plugins, with technical input from A.G, and conceptual inputs from A.G, S.T, L.G, V.D-E. A.G, J.V, and V.D-E analyzed the data, with inputs from S.T, L.G, P.R and P-F.L. S.T, V.D-E, A.G and J.V wrote the manuscript with inputs from L.G, P-F.L and P.R. All authors discussed the scientific results and their interpretation. A.G, V.D-E, J.V and L.G created the figures and visualizations, P-F.L, S.T, L.G and P.R acquired funding.

## ABBREVIATIONS

SNR: signal-to-noise-ratio
IoU: intersection over union
ph3: phospho-histone H3
PBS: phosphate-buffered saline
FRC-QE: Fourier ring correlation quality estimate
Ecad: E-cadherin
H2B: Histone 2B

## V. SUPPLEMENTARY INFORMATION

### A. Supplementary Movies

- **Suppl.Movie 1**: 3D rendering of a 78 h small sample processed through the pipeline. Hoechst is in grey, T-Bra in magenta, Oct4 in green, and FoxA2 in blue.
- **Suppl.Movie 2**: Z-stack of a 78 h big sample processed through the pipeline. Hoechst in grey, T-Bra in red, Oct4 in green, and FoxA2 in blue. Depth in the sample is indicated in the bottom right corner.
- **Suppl.Movie 3**: Z-stacks of 8 samples imaged at 78 h all in the same acquisition, and processed through the pipeline. Hoechst in gray, T-Bra in red, Oct4 in green, and FoxA2 in blue. Depth in the sample is indicated in the bottom right corner.
- **Suppl.Movie 4**: Z-stacks of two samples stained with ph3, on the right at 72 h and on the left at 96 h. Detected divisions are visualized as multicolor ROIs superimposed with the image, where only the contour is shown.

### B. Supplementary Figures

**Fig. S1.**
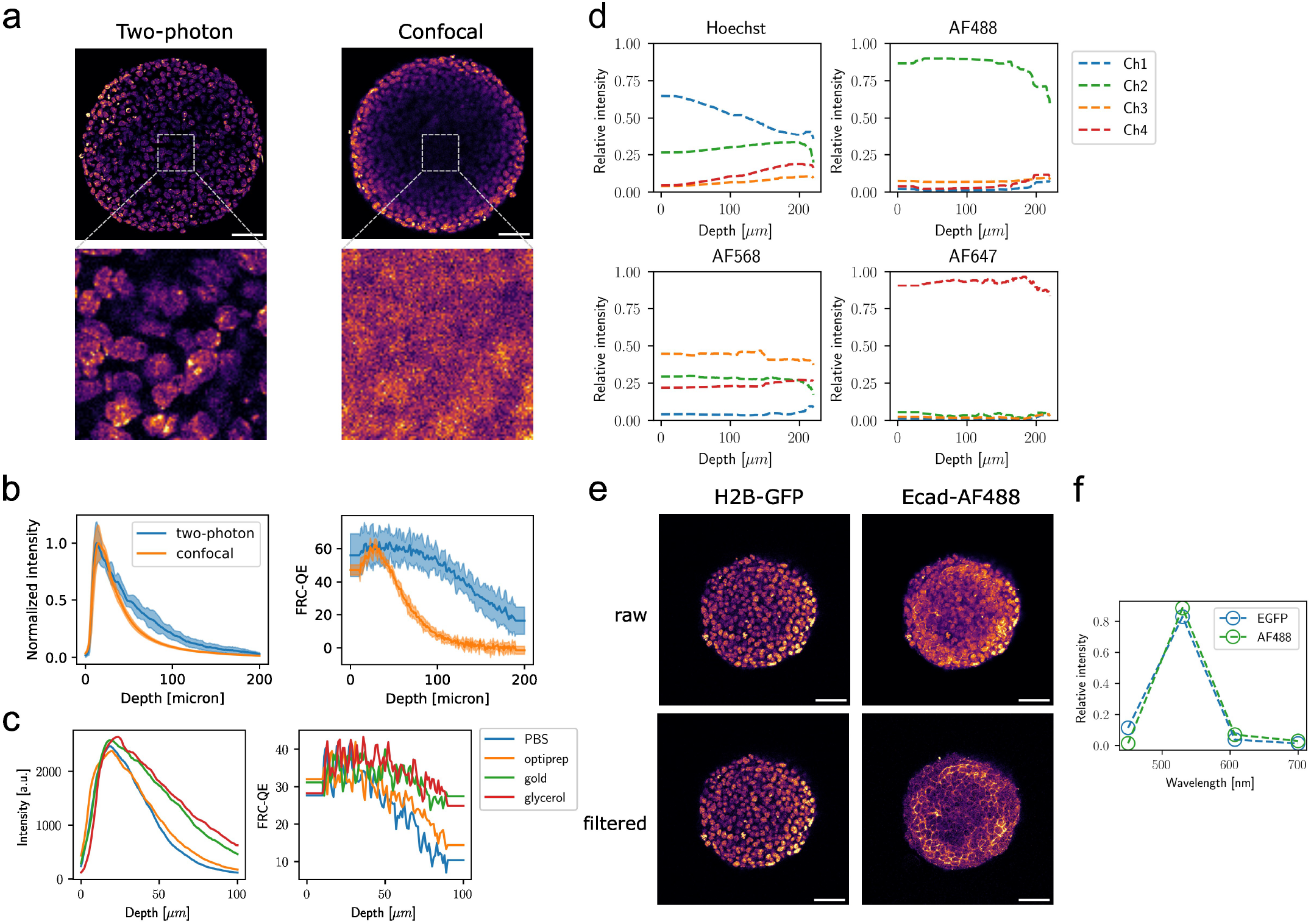
Benchmarking of dual-view two-photon imaging and spectral unmixing. **(a)** Two-photon and confocal images of Hoechst-stained 77 h gastruloids at 100 *µm* depth. Insets show nuclei in the central 100×100 pixels. Scale bars are 50 *µm*. **(b)** Normalized intensity and FRC-QE in the center of gastruloids described in (a), as a function of depth. Shaded regions show standard deviation across six gastruloids of similar size. **(c)** Normalized intensity and FRC-QE obtained with two-photon imaging in the center (60 × 60 *µm*^2^ region) of 72 h Hoechst-stained gastruloids as a function of depth, mounted in either PBS, 20% optiprep, ProLong gold antifade, or 80% glycerol medium. **(d)** Normalized relative intensities detected in four detector channels as a function of imaging depth on aggregates stained with either Hoechst, AF488, AF568, or AF647. **(e**,**f)** Spectral unmixing of images acquired on gastruloids expressing transgenic H2B-GFP, additionally stained with Ecad-AF488. Raw data show significant cross-talk, particularly in the second channel. Unmixing using the spectra shown in (f) strongly reduces nuclei cross-talk in the Ecad-AF488 channel.

**Fig. S2.**
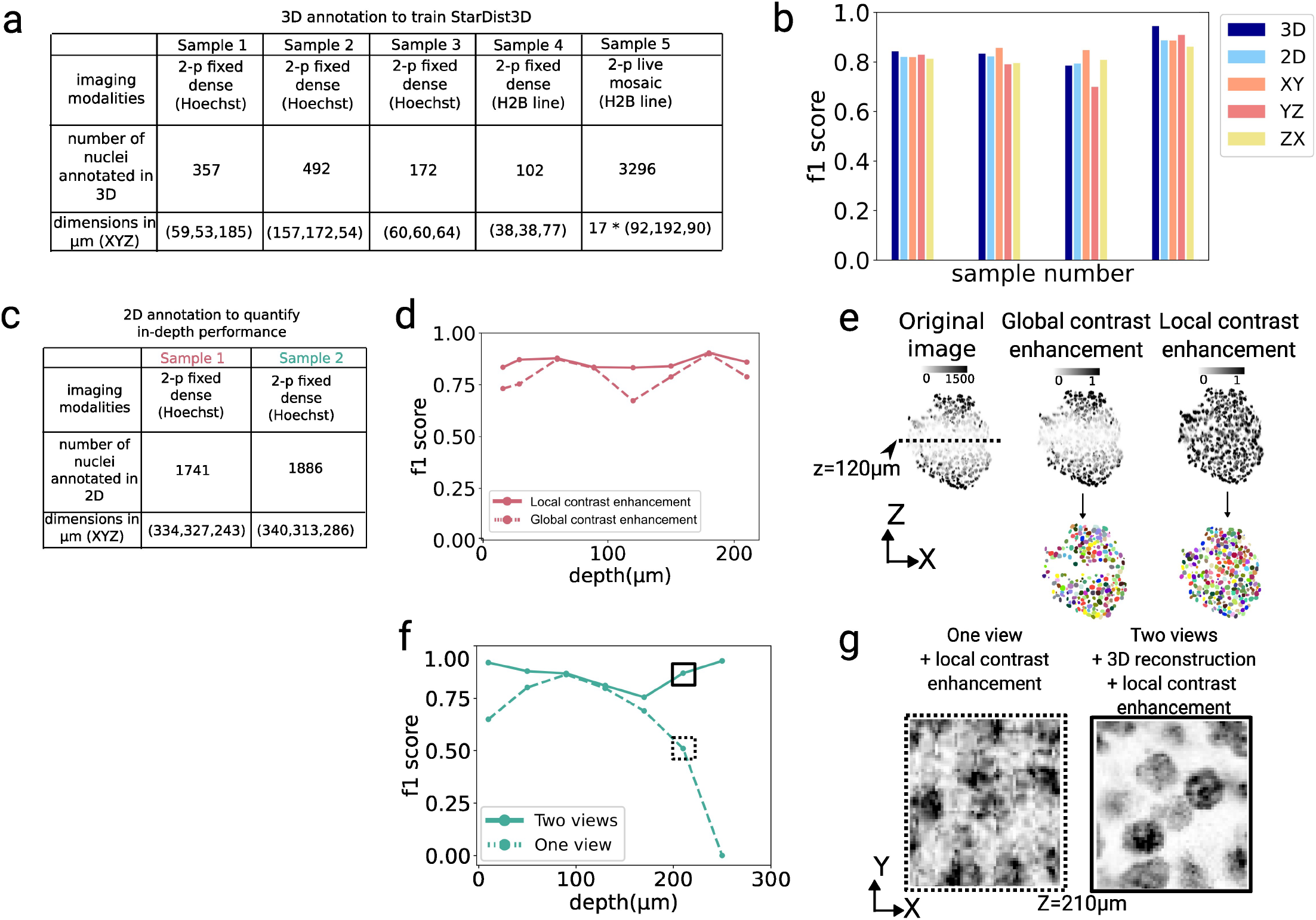
Quantification of StarDist3D performance. **(a)** Training dataset of StarDist3D. Five samples with different characteristics were annotated in 3D, all the nuclei in a cropped region, with dimensions indicated in the table. (**b**) Comparison for each sample of the F1 scores obtained using either 3D or 2D ground-truths. 2D annotations were obtained on XY slices sampled every 20 *µm*. For each slice, we quantified the F1 score at IoU threshold = 0.5. To check for eventual biases due to the original anisotropy of the image, we also plot the F1 scores using 2D annotations in YZ and XZ (yellow and pink bars). Light blue bars represent the average of scores along all 2D planes. The F1 score using 3D annotations (dark blue bars) is well approximated by the F1 score using 2D annotations in XY, with a 5% difference at most. (**c**) As a consequence, we could quantify the segmentation score on new samples only by annotating in 2D the XY planes of two samples. 7 to 8 slices were annotated regularly throughout the volumes and the plots (**d-g**) show the 2D quantification of nuclei segmentation performance in depth. (**d**) F1 score as a function of depth in sample 1, comparing our contrast enhancement method in a local window (solid line) versus using a classical contrast enhancement method based on histogram statistics from the full image (dotted line), see Methods. The arrow shows the midplane, where the SNR is the lowest after fusion of both sides. (**e**) XZ view of the images and corresponding segmentation using either global or local contrast enhancement methods. The dotted line indicates the midplane. (**f**) F1 score as a function of depth in sample 2, comparing two-views imaging (solid line) versus one view imaging on the same sample (dotted line). We enhanced contrast locally in both samples to increase the signal in deeper z planes, but two-views imaging is still necessary to segment the nuclei deeper than 200 *µm* in the sample. (**g**) Cropped regions at 210 *µm* depth of a sample imaged from two sides, registered, reconstructed in 3D, and locally enhanced, versus the same image from only one side, far from the objective.

**Fig. S3.**
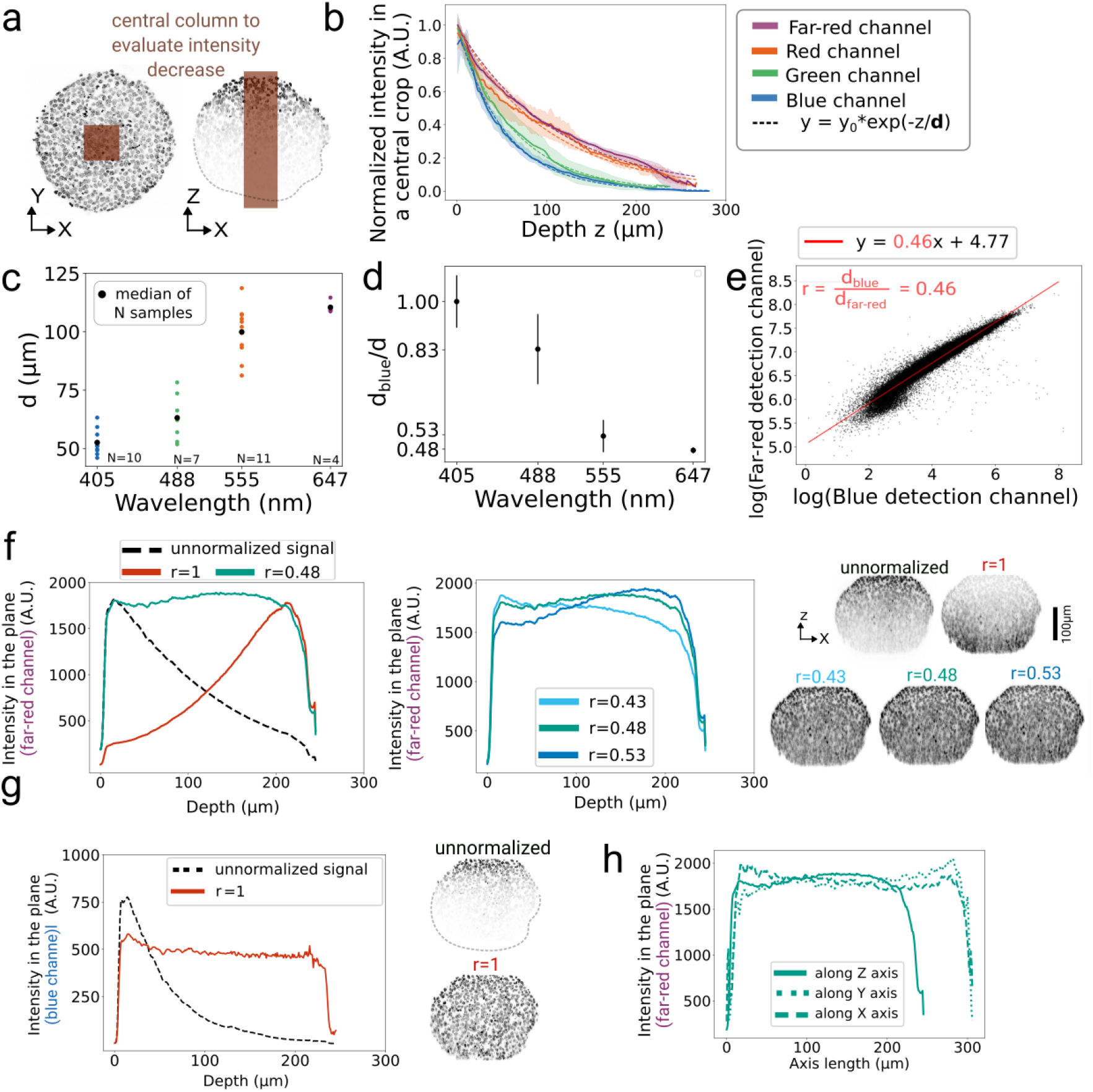
The wavelength-dependent normalization method. **(a)** We evaluated how the intensity decreases in an ubiquitous signal as a function of depth and wavelength. Samples are immunostained with histones markers in the four colors. The intensity is measured in a central column of the sample, to simplify the geometry and remove border effects. In this region, we compute the median intensity in each plane. **(b)** Plot of the intensity in each channel, normalized to 1. Shaded regions indicate standard deviation. An exponential function is fitted to the normalized intensity to extract the characteristic length of decrease d. **(c)** This length d is plotted for each sample as a function of their wavelength. **(d)** We extract the ratio *r* = *d*_*blue*_*/d, d*_*blue*_ being the median of the characteristic length d in the blue channel. Mean and standard deviation are shown. **(e)** Plot of the intensity in the far-red channel as a function of the intensity in the blue channel, both axis are in logarithmic scale. Images are binned so that each black dot represents a 10 · 10 *µm*^2^ square. N=5 samples are stacked onto the plot.The slope of 0.46 validates the value 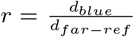 **(f)** Using the method described in Fig. S8c, we measure the intensity in the far-red channel as a function of depth, using the local normalization method described in the Methods III J, and different values of the ratio *r* = *d*_*blue*_*/d*_*far*−*red*_. The black dotted line is the non-normalized signal, the red line represents the case *r* = 1 where we apply the correction without taking into account the wavelength-dependency of the intensity decrease. The blue line represents the ratio found experimentally and cyan and blue lines show the profile for close values. Corresponding XZ views are shown on the right. **(g)** Similarly to panel (e), intensity in depth for the blue channel, without normalization and using r=1, which flattens the profile in depth as shown also in the XZ views. **(h)** Using the experimental value *r* = 0.48, we show for the far-red channel the profile obtained plane by plane by slicing either in Z, X or Y. The flat profiles shows no big heterogeneities in the normalized sample.

**Fig. S4.**
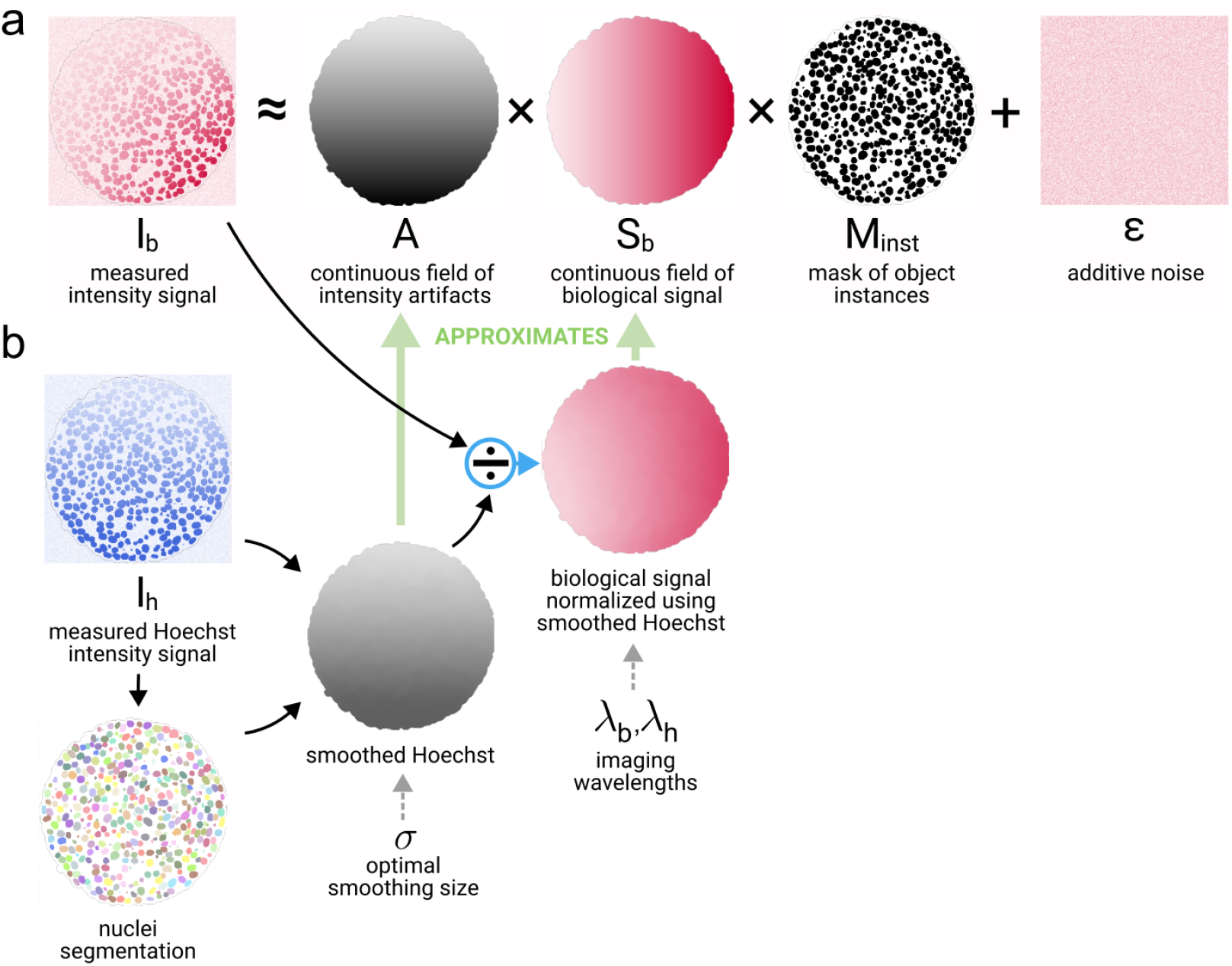
Illustration of the intensity model and of the normalization procedure based on a ubiquitous signal. (a) In equation (9), we model any arbitrary intensity signal *I*_*b*_ by decomposing it into three terms: (i) a continuous field *A* representing local intensity artifacts, (ii) a field *S*_*b*_ representing a continuous depiction of the biological signal, and (iii) an instance mask *Minst*, which localizes the biological objects (e.g. nuclei or membranes) from where the biological signal originates. The first two fields are assumed to have variations at length scales equal or above the typical cell size. Finally, an additive term *ε* which varies at sub-cellular length-scales and encompasses all the remaining voxel-scale signal (e.g. measurement noise or intra-cellular gene expression heterogeneities) completes the model. **(b)** An ubiquitous signal (here Hoechst) can be used to approximate the field of intensity artifacts, which can be removed from the original intensity signal.

**Fig. S5.**
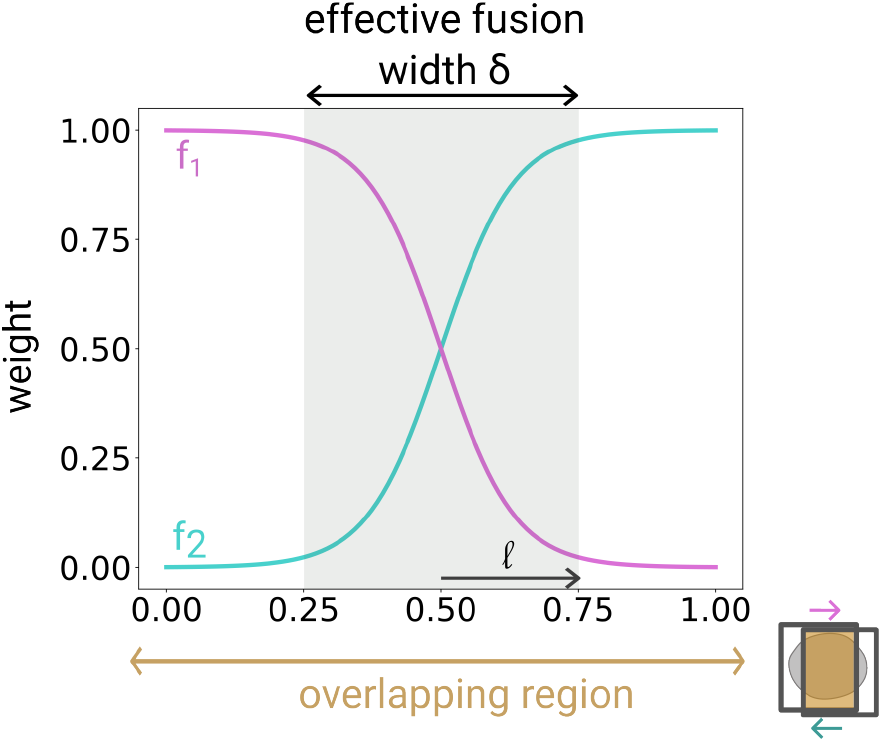
Illustration of the sigmoid function that is used for weighted fusion of dual-view images, see equation (3). The coordinate along the overlapping region between the two views is mapped to the range [0,1] to define the parameters of the sigmoid function independently from the gastruloid size. *l* is the decay length of the sigmoid, and we define the effective fusion width δ as the fixed fraction of the overlapping region where both signals are substantially present to create the fused part.

**Fig. S6.**
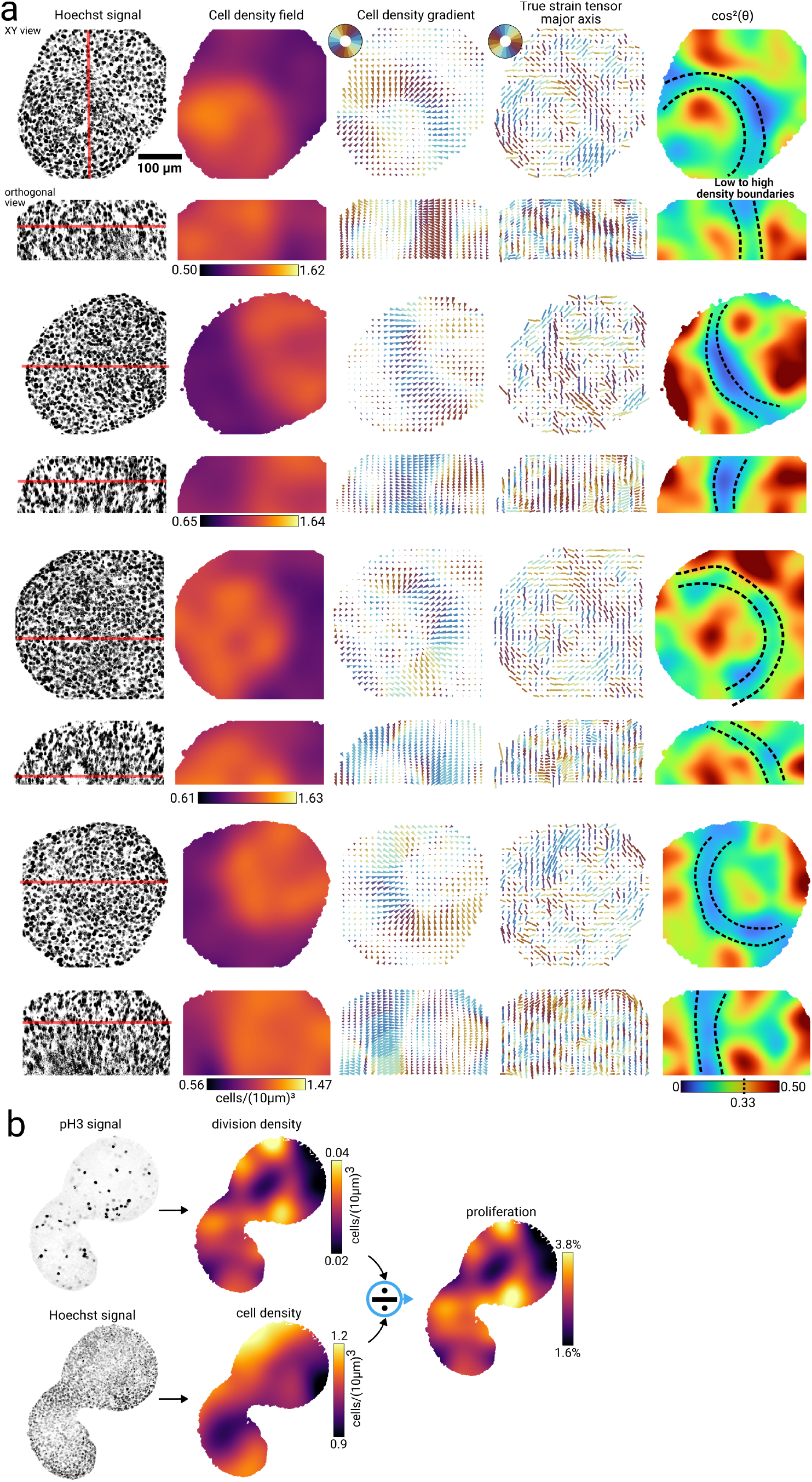
**(a)** Illustration of the reproducibility of the spatial colocalization of a sharp low-to-high density boundary with a region of high nuclei alignment in XY and orthogonal views from several gastruloids. The leftmost column shows locally enhanced Hoechst signal in the XY plane, and a plane perpendicular to it. The red lines give the positions of the complementary plane. 120 h gastruloid, stained with ph3 and Hoechst, processed and segmented (nuclei and divisions) using the pipeline to generate the division density map and the cell density map. With these two fields, we compute a 3D proliferation map (shown here is a 2D section in the mid-plane).

**Fig. S7.**
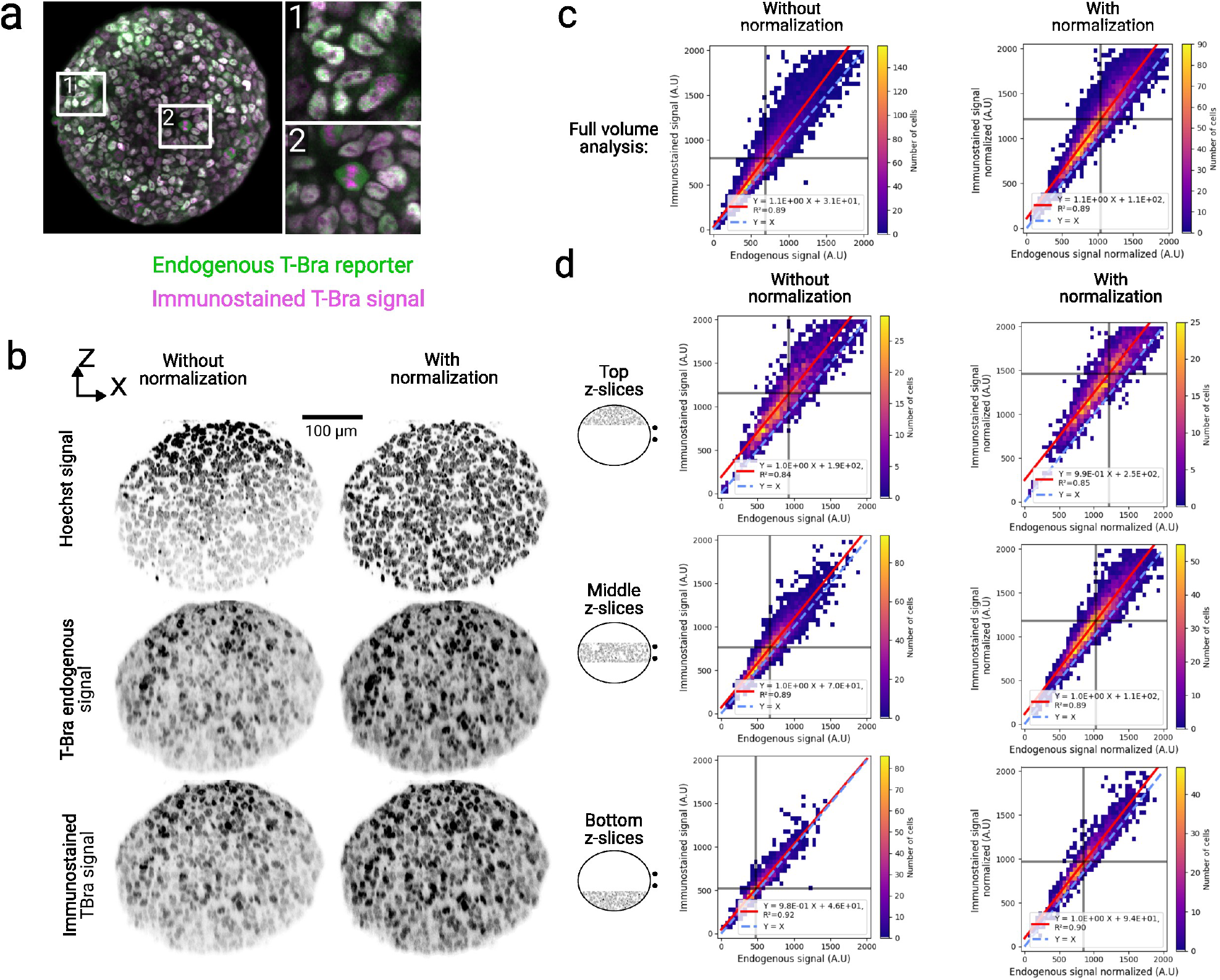
Direct comparison of depth-dependent correlation of intensity between immunostained T-Bra signal and the intensity signal of a T-Bra reporter (H2B-GFP expressed under a T-Bra promoter) in gastruloids grown from a transgenic cell line fixed at 72 h. **(a)** Image of a sample expressing the endogenous T-Bra reporter (green) and immunostained T-Bra (magenta). Signals are correlated almost everywhere (1) except for cells in mitosis (2) **(b)** Comparison of XZ views of Hoechst, endogenous T-Bra and immunostained T-Bra, before and after wavelength-dependent intensity normalization using the Hoechst signal as reference. **(c)** Intensity correlation heatmaps between the immunostained and endogenous T-Bra signals, in the full sample. After normalization, the two signals are directly equal up to a small additive bias. Vertical and horizontal black lines represent the median of each signal. **(d)** Intensity correlation heatmaps computed in three 3D slices of width 50 *µm* at varying depths. Normalization leads to a totally homogeneous correlation relationship between the two signals, with the same slope and additive bias at all depths.

**Fig. S8.**
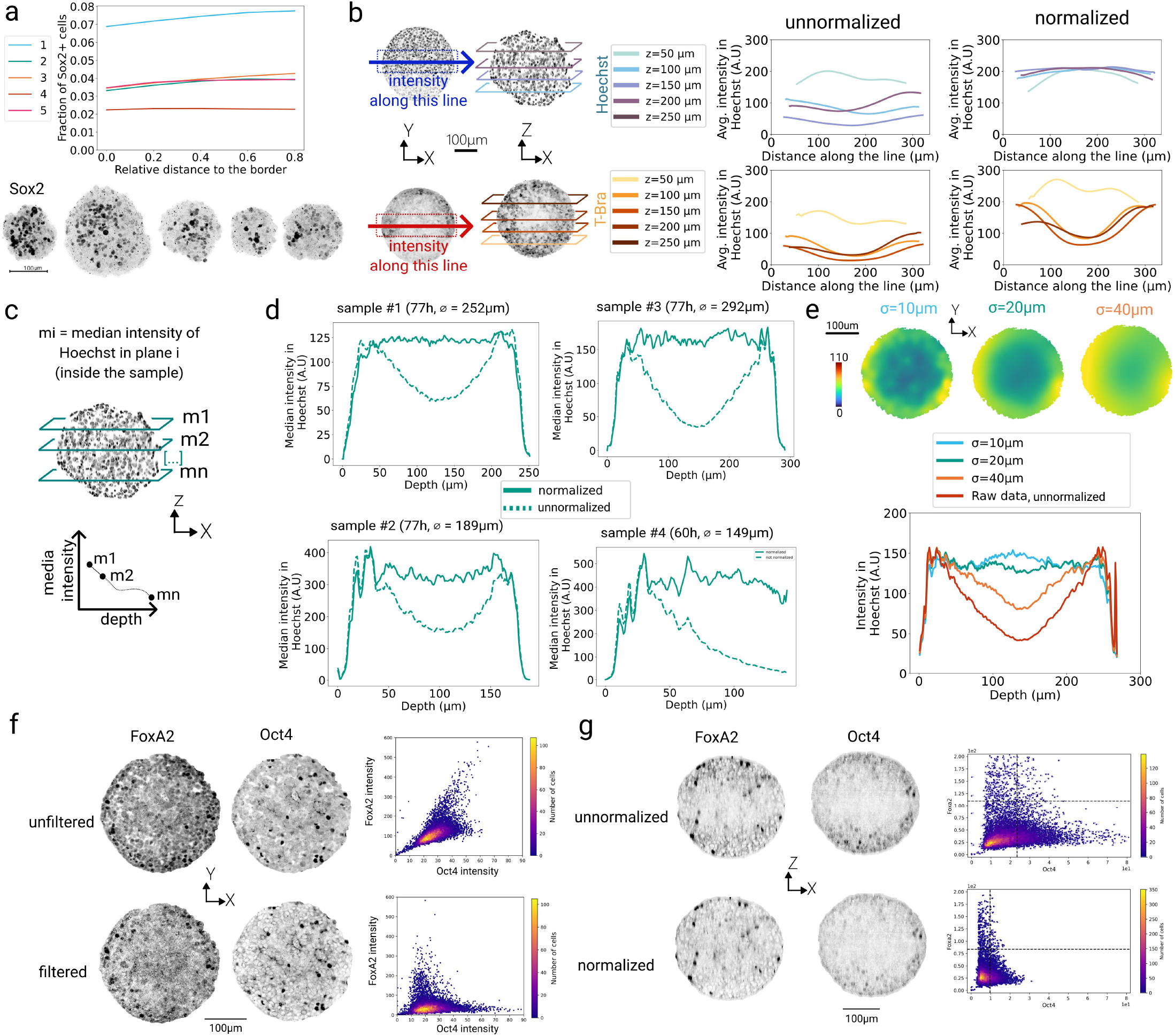
Effect of spectral filtering and local intensity normalization on 3D gene expression analysis. **(a)** As illustrated in Fig.5d for FoxA2, we analyze the radial distribution of Sox2 in 60 h gastruloids. Below, we present 3D visualizations of the normalized Sox2 signal alongside the segmentation of Sox2+ cells. The resulting 3D maps of the local fraction of Sox2+ cells reveal no preferential radial pattern. **(b)** Comparison of lateral intensity profiles at different depths. We compute the intensity across rectangular sections of each image in 2D, as shown on the left. The graphs show the line profiles at different depths for the Hoechst channel (top) and the T-Bra channel (bottom), comparing profiles before (left plot) and after (right plot) normalization. Color code stands for depth (darker colors correspond to deeper z-sections). We observe for non-normalized data that z-planes at mid-depth have a low intensity that is corrected in the normalized case. Hoechst profiles are flattened throughout the section after normalization, as expected. **(c)** Illustration of plots (d) and (e) : for each z-plane, we measure the median intensity in the Hoechst channel, within the sample, and plot it as a function of its depth. We use this method to quantitatively check that the intensity in the nuclei channel is constant in depth and does not present gradients after normalization. **(d)** Using the method described in (c), we show the median intensity as a function of depth for 4 samples from different acquisitions, corresponding to different timing and/or diameters. We compare the intensities before (dotted lines) and after normalization (solid lines), and observe that the normalization scheme robustly flattens the intensity profile. **(e)** For one sample, we compare different values of *σ* which is the standard deviation of the Gaussian kernel used to normalize the data. The images are respective maps of nuclei intensities for *σ* = 10, 20 and 40 *µm*. Using the method described in (d), we show the median nuclei intensity as a function of depth for these three *σ* values. **f** Correlation maps between FoxA2 and Oct4 for spectrally non-filtered data (top) versus filtered data (bottom), and the corresponding images shown on the left. Besides filtering, images were still registered and normalized the same way. **(g)** Correlation maps between FoxA2 and T-Bra for non-normalized data (top) versus normalized data (bottom), and the corresponding images are shown on the left. Images were still filtered and registered the same way.

**Fig. S9.**
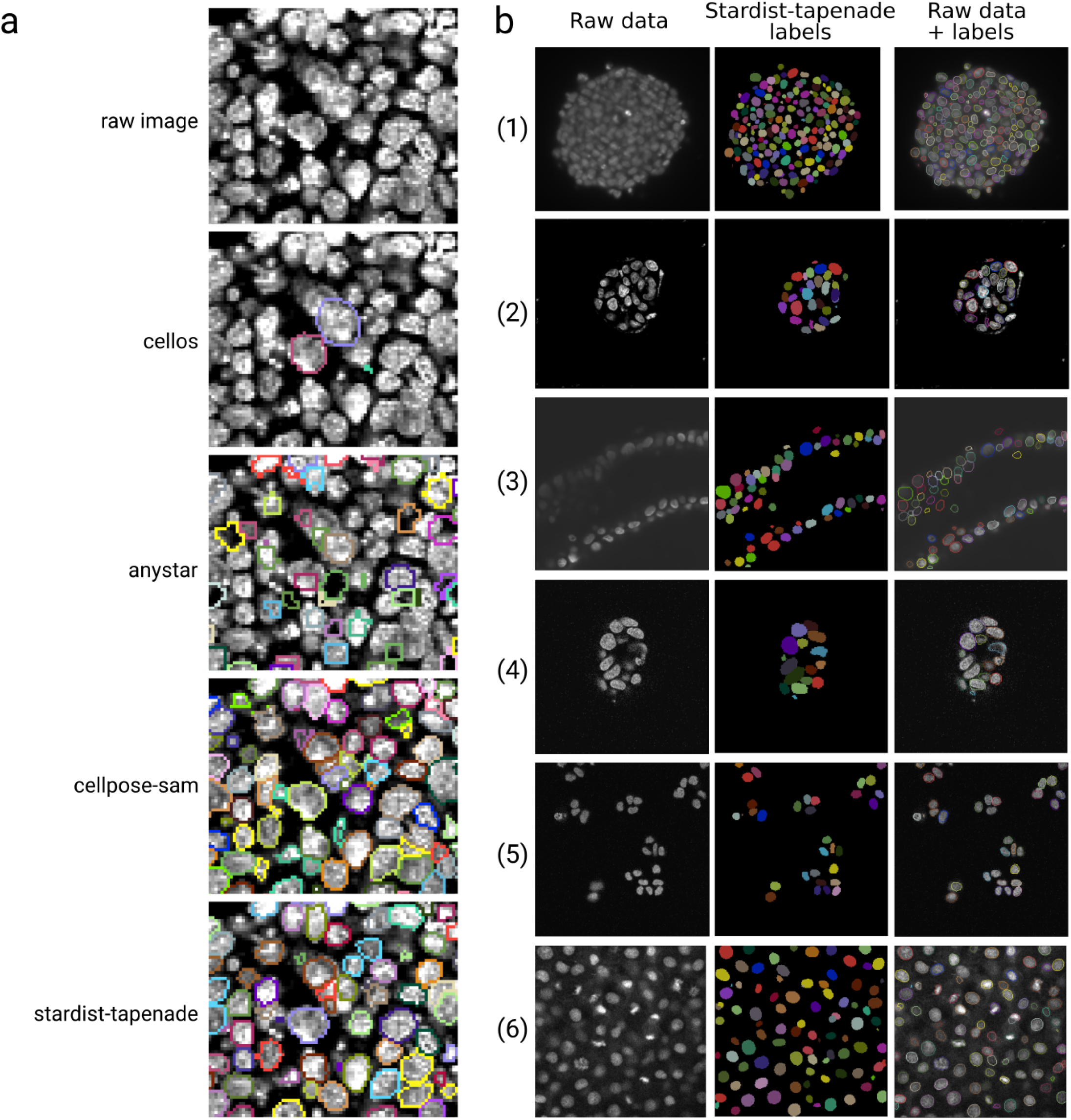
Qualitative benchmark of our segmentation model. All images were segmented in 3D, and we show here one XY plane for simplicity. **(a)** Qualitative assessment of the performance of different segmentation models against stardist-tapenade on gastruloid images (Hoechst staining). The pre-trained Cellpose-SAM [70] model gives high-quality results, which are further investigated in Fig. S10. **(b)** Qualitative comparison of our custom Stardist3D segmentation strategy (stardist-tapenade) on diverse published 3D nuclei datasets proves the versatility of our model. (1) Gastruloid stained with the nuclear marker DRAQ5 imaged with an open-top dual-view and dual-illumination LSM [62]. (2) Breast cancer spheroid [29]. (3) Primary pancreatic ductal adenocarcinoma organoids imaged with confocal microscopy [29]. (4) Human colon organoid imaged with LSM laser scanning confocal microscope [29]. (5) Monolayer HCT116 cells imaged with LSM laser scanning confocal microscope [29]. (6) Fixed zebrafish embryo stained for nuclei and imaged with a Zeiss LSM 880 confocal microscopy [63].

**Fig. S10.**
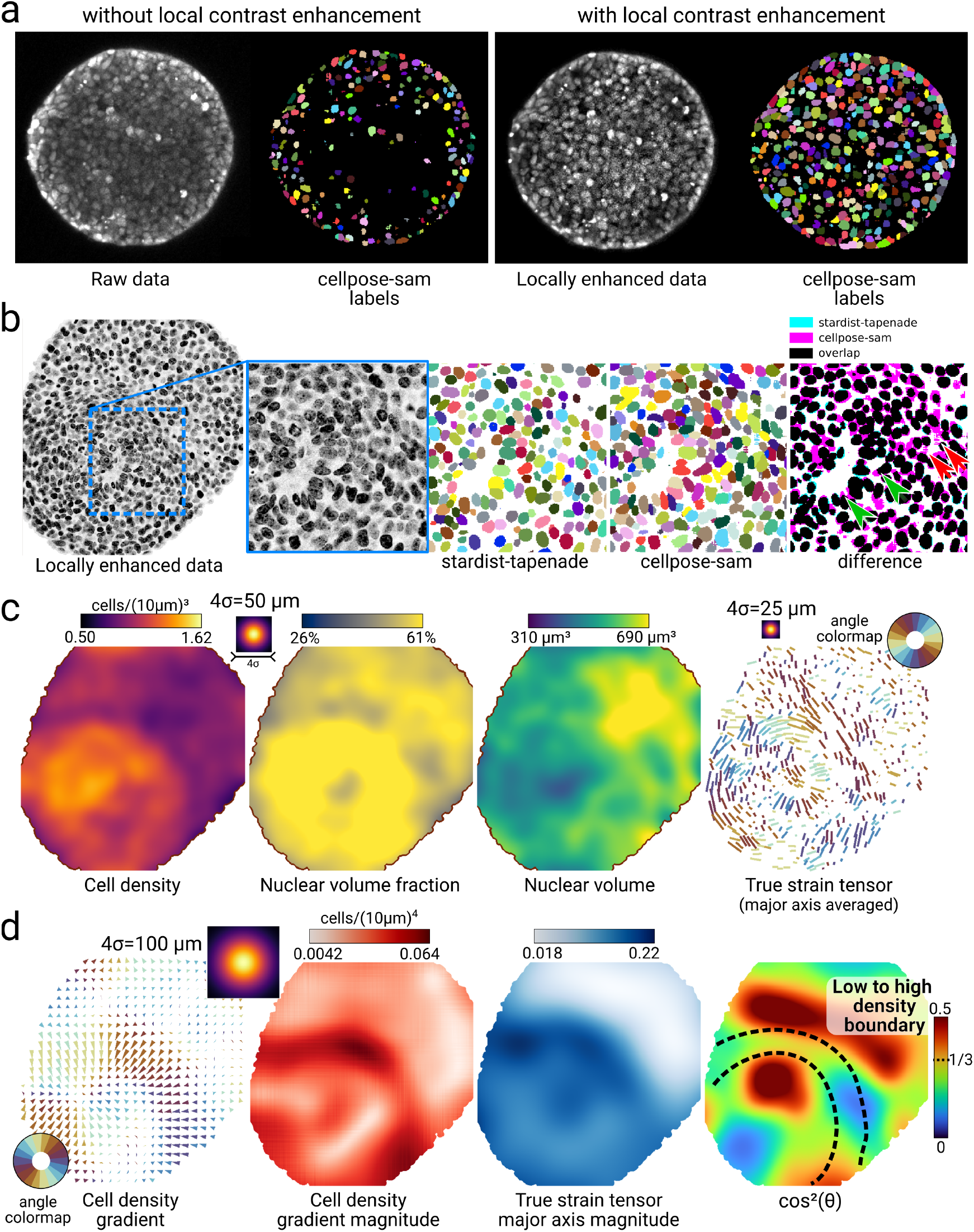
Validation of the Cellpose–SAM pretrained model for nuclear segmentation and morphology in gastruloids. **(a)** The Tapenade preprocessing pipeline improves Cellpose–SAM [70] outputs, especially in deep planes (*z* > 150 *µ*m), by using local contrast enhancement. **(b)** Compared to our StarDist3D model (stardist-tapenade), Cellpose–SAM better preserves elongated nuclei (green arrows), while StarDist3D tends to under-segment by rounding objects. Cellpose–SAM can, however, yield partial segments in low–signal-to-noise regions (red arrows). Overall this model seems highly competent for nuclei segmentation in gastruloids. Here and in the rest of this figure, we used the same sample as in Fig. 4 to allow for direct comparison. **(c)** Tissue-scale, packing-related metrics from Cellpose–SAM labels qualitatively match those from stardist-tapenade, though the more accurate boundaries increase estimated nuclear volume and volume fraction. The true-strain major-axis field remain similar because StarDist3D mostly preserves orientations. **(d)** Quantification of the region where the cell-density gradient and the true-strain major axes are anti-aligned is consistent when using Cellpose–SAM. Vector magnitudes of both fields are displayed to aid interpretation in areas with small gradients or deformations.

